# Ecological basis and genetic architecture of crypsis polymorphism in the desert clicker grasshopper (*Ligurotettix coquilletti*)

**DOI:** 10.1101/2021.04.29.441881

**Authors:** Timothy K. O’Connor, Marissa C. Sandoval, Jiarui Wang, Jacob C. Hans, Risa Takenaka, Myron Child, Noah K. Whiteman

**Author notes:** **Author contributions:** TKO and NKW designed research. TKO, JCH, RT, and MC collected specimens and surveyed predation environments. TKO, JW, and MCS extracted DNA, and TKO and JW prepared sequencing libraries. TKO and MCS prepared image data. TKO performed analyses. TKO wrote the manuscript with input from all authors.

## Abstract

Color polymorphic species can offer exceptional insight into the ecology and genetics of adaptation. Although the genetic architecture of animal coloration is diverse, many color polymorphisms are associated with large structural variants and maintained by biotic interactions. Grasshoppers are exceptionally polymorphic in both color and karyotype, making them excellent models for understanding the ecological drivers and genetic underpinnings of color variation. Banded and uniform morphs of the desert clicker grasshopper (*Ligurotettix coquilletti*) are found across the western deserts of North America. To address the hypothesis that predation maintains local color polymorphism and shapes regional crypsis variation, we surveyed morph frequencies and tested for covariation with two predation environments. Morphs coexisted at intermediate frequencies at most sites, consistent with local balancing selection. Morph frequencies covaried with the appearance of desert substrate – an environment used only by females – indicating that ground-foraging predators are major agents of selection on crypsis. We next addressed the hypothesized link between morph variation and genome structure. To do so, we designed an approach for detecting inversions and indels using only RADseq data. The banded morph was perfectly correlated with a large putative indel. Remarkably, indel dominance differed among populations, a rare example of dominance evolution in nature.

## INTRODUCTION

Animal coloration has manifold ecological roles with profound effects on fitness (Caro 2005). Therefore, species with variable coloration provide excellent windows into the ecological drivers and genetic basis of adaptation. Although the study of animal color dates to the founding of evolutionary biology (Darwin 1859; Wallace 1877), color polymorphic species continue to yield new insight into general evolutionary processes (Gray and McKinnon 2007; Forsman et al. 2008; Svensson 2017; Orteu and Jiggins 2020). Crypsis polymorphisms (the coexistence of discrete camouflage morphs) have been particularly important in illuminating how natural selection maintains adaptive genetic variation within populations (balancing selection *sensu lato*). One possibility is selection in the face of maladaptive gene flow (migration-selection balance), which can operate both regionally (e.g., Hoekstra et al. 2004; Rosenblum 2006) and locally (i.e. Levene’s model of spatially varying selection (1953); Sandoval 1994b). Other local processes include negative frequency-dependent selection, which can arise from a variety of processes including predator foraging behavior (Bond 2007), and sexually antagonistic selection driven by contrasting fitness consequences of color morph between sexes (Forsman 1995; Bonduriansky and Chenoweth 2009).

While the adaptive value of coloration may appear self-evident, it should not be assumed (Jones et al. 1977). Correlations between an organisms phenotype and its environment can provide *prima facie* evidence that natural selection acts on color polymorphism (Endler 1977, 1986) and point to processes that might maintain phenotypic diversity. If most populations are nearly monomorphic, this suggests that maladaptive gene flow or recurrent mutation oppose local adaptation. If instead most populations have intermediate morph frequencies, local processes such as negative frequency dependence or fine-scale spatially varying selection are implicated. Covariation of morph

frequencies and environmental variables suggest the local environment determines the polymorphic equilibrium, and the particular variables correlated to morph frequencies can identify plausible selective agents.

While classic studies revealed that color polymorphisms are often Mendelian traits, recent work has shown that similar inheritance patterns can belie a diversity of genetic bases (San-Jose and Roulin 2017; Orteu and Jiggins 2020). Coding SNPs (Theron et al. 2001; Nachman et al. 2003; Mundy et al. 2004; Rosenblum et al. 2004; Cooke et al. 2017), transposable element insertions (van t’Hof et al. 2016; Woronik et al. 2019), and cis-regulatory polymorphisms (Lewis et al. 2019; Tian et al. 2019) have all been implicated in within-species color variation.

Notably, color polymorphisms are often associated with structural variants, including chromosomal inversions and large insertions or deletions (indels) (Wellenreuther and Bernatchez 2018). By locking alternative allele combinations into non-recombining haplotypes, inversions can create “supergenes” with simple inheritance but potentially complex phenotypic effects (Thompson and Jiggins 2014). Inversions have been associated with polymorphic mimicry in butterflies (Joron et al. 2011; Kunte et al. 2014; Nishikawa et al. 2015), plumage variation in birds (Thomas et al. 2008; Küpper et al. 2015; Lamichhaney et al. 2015; Tuttle et al. 2016), and crypsis in stick insects (Lindtke et al. 2017). Indels have also been linked to color variation, although in fewer taxa (e.g., Gallant et al. 2014). A green-brown polymorphism in *Timema* stick insects was recently mapped to a ∼5 Mb deletion that spans loci underlying continuous color variation in related species (Villoutreix et al. 2020). Analogous to an inversion, a large indel can therefore convert a polygenic trait into a discrete dimorphism (Gutiérrez-Valencia et al. 2021).

Irrespective of color variation’s genetic basis, the dynamics of selection depend upon allelic dominance at causative loci (Manceau et al. 2010; Rosenblum et al. 2010; Nuismer et al. 2012; Laurent et al. 2016). The visibility of each allele to natural selection affects the probability of stochastic loss, the rate of phenotypic change, and the equilibrium allele frequencies under an array of selective regimes. Characterizing allelic dominance is therefore essential to understanding adaptation in natural populations.

Grasshoppers and their relatives (Orthoptera, infraorder Acrididea) are important models in both evolutionary ecology and genetics (Haldane 1920; Nabours et al. 1933; Fisher 1939; White 1973). Like other orthopterans, grasshoppers are notable for the extent of polymorphism in both color (Rowell 1972; Dearn 1990) and karyotype (White 1973; Bidau and Martí 2010). They are therefore an excellent group with which to study the links among balancing selection, color polymorphism, and structural variation. The desert clicker (Acrididae: *Ligurotettix coquilletti*) is a cryptic grasshopper that is ubiquitous throughout the Sonoran, Mojave, and Peninsular Deserts, with isolated populations in the Great Basin (**Fig 1A**). Two discrete color morphs occur in both sexes (McNeill 1897; Rehn 1923): a uniform morph with relatively homogeneous color, and a banded morph with contrasting light and dark bands along the body axis (**Fig 1B**). Although both morphs sometimes cooccur (Rehn 1923; Chapman 1991), their geographic distribution and frequency have not been surveyed.

**Figure 1.**
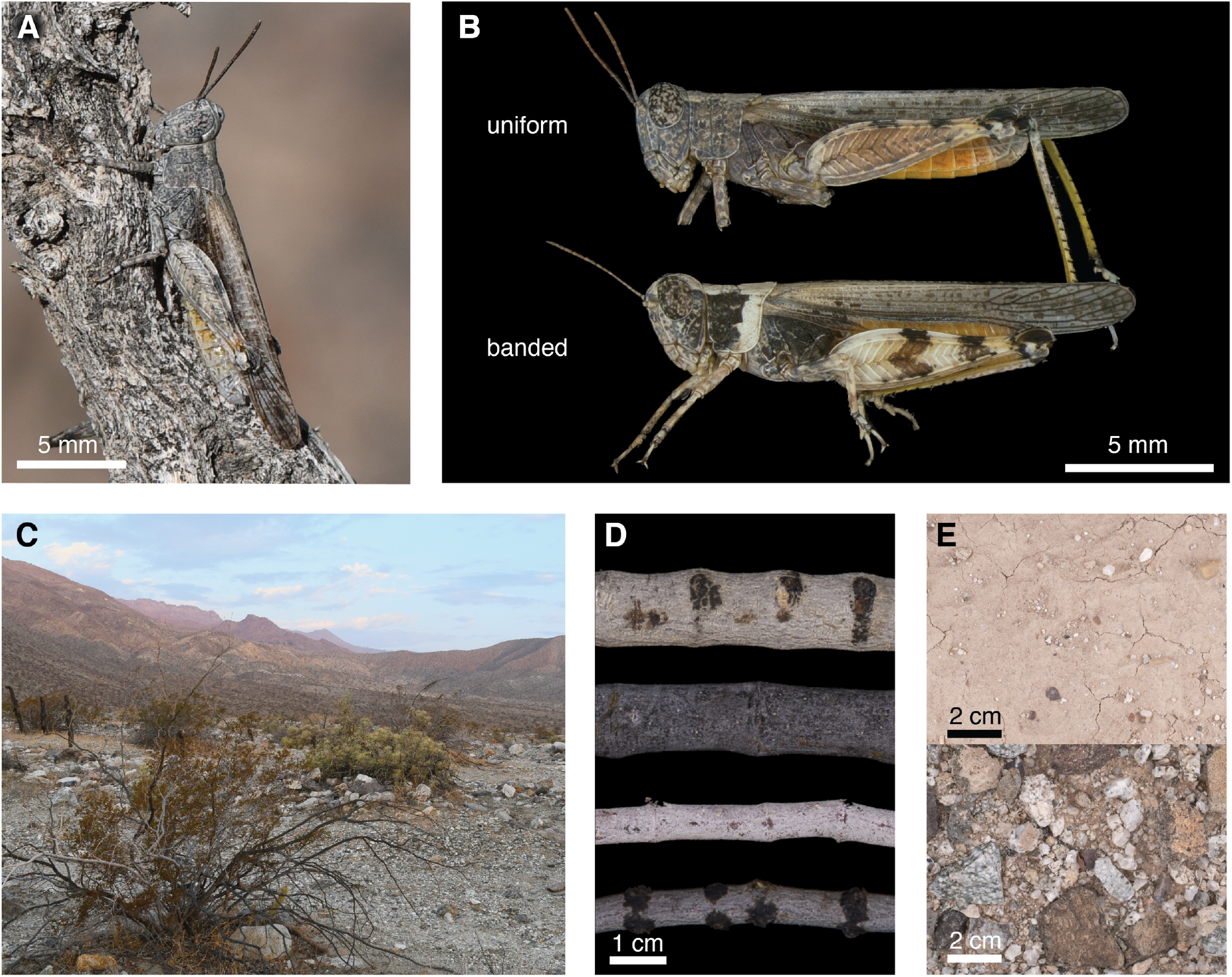
**A**. The desert clicker grasshopper, *Ligurotettix coquilletti* on greasewood (*Sarcobatus vermiculatus*; Mina, NV). **B**. Two discrete color morphs of the desert clicker (Ocotillo Wells, CA). **C**. Typical desert clicker habitat, with creosote bush (*Larrea tridentata*) in foreground (Boyd Deep Canyon Desert Research Station, Palm Desert, CA). **D**. Both sexes spend most of the day on host plant stems (primarily creosote bush), which vary in color and patterning. **E**. Adult females spend several hours each morning ovipositing into desert substrate, ranging from low-contrast silt (top) to highly-patterned granitic gravel (bottom).

The life history of desert clickers is tightly associated with their host plants. Most populations are found on creosote bush (Zygophyllaceae: *Larrea tridentata*), although saltbrush (Chenopodiaceae: *Atriplex spp*.) and greasewood (Sarcobataceae: *Sarcobatus vermiculatus*) are the primary host plants in the Great Basin. After hatching from an egg embedded in rocky substrate, males quickly move to a host plant and virtually never return to the ground (Wang and Greenfield 1994; **Fig 1C**): “It is as truly bush-loving as any acridid of my acquaintance…” (Rehn 1923). Males call insistently from plant stems (**Fig 1D**) for up to 16 hours per day (Greenfield 1992), the sound and movement of which may expose their position to predators. Robber flies (Diptera: Asilidae), spiders, mantids, *Capnobotes* spp. katydids, and lizards (*Uta* spp. and *Cnemidophorus* spp.) depredate desert clickers on host plants, and cactus wrens (*Campylorhynchus brunneicapillus*) are also likely predators (M. Greenfield, pers. comm.). Females also perch, evaluate mates, and copulate within host plants throughout most of the day. However, adult females also descend to the desert substrate for 3–4 hours spanning dawn (Wang 1990, cited in Greenfield 1992, **Fig 1E**). A portion of this time is spent ovipositing, during which females are immobilized for up to 45 minutes (Wang and Greenfield 1994). Many desert vertebrates are most active at dawn (e.g., Ricklefs and Hainsworth 1968; Baumgardner et al. 1980; Huey 1982). Therefore, females may be particularly vulnerable to predation during oviposition. Whether predators select for different color morphs on stem vs. substrate – and whether such selection varies across the species’ range – is unknown.

Here we report results of our study on the ecological and genetic basis of crypsis polymorphism in desert clickers. We first addressed the hypothesis that balancing selection maintains adaptive color variation within populations. To do so, we first collected desert clickers across the entire species’ range and estimated morph frequencies at 20 focal sites. We then tested for correlations between morph frequencies and variation in two predation environments (plant stems and desert substrate). With these correlations we identified processes that may maintain polymorphism within populations and underlie morph frequency differences among populations.

Second, we hypothesized that color variation in desert clickers is associated with large-scale structural variation. As grasshoppers are non-model taxa with large and repetitive genomes (Wang et al. 2014; Verlinden et al. 2020), we lacked the resources to exhaustively describe structural variants (Ho et al. 2020). Instead, we designed and implemented a new approach to detect structural variants that distinguish two phenotypic classes using only RADseq data. In parallel, we performed an *F*ST outlier scan to identify loci associated with color morph. With results of these phenotype-genotype correlations we characterized dominance relationships of color-associated loci and tested for molecular genetic evidence of balancing selection.

## METHODS

### Survey of grasshopper phenotypes and environmental variation

#### Phenotype survey

To estimate the frequency of desert clicker morphs across populations, we surveyed 20 sites in Arizona, California, and Nevada in August–September, 2018 (**Fig S1**.**1, Table S1**.**1**). At each site we collected grasshoppers (*N* = 312, range: 4–25 per site) and noted the color morph of individuals that we observed but did not capture (*N* = 165; range: 0–24 per site) for a total of 488 phenotype observations (range: 4–48 per site). We further analyzed the 19 sites with ≥ 10 observations.

#### Photography

We photographed live, chill-immobilized grasshoppers in lateral view to record phenotypes. From each site we also collected and photographed six 15-cm stem segments (three 6 mm in diameter and three 12 mm in diameter) from each of ten host plants (typically creosote bush, but saltbrush and greasewood at one site each; see **Fig S1**.**1**). These stem sizes span the typical size range of stems where desert clickers perch. We photographed bare substrate 1.5m away from host plants to capture the visual environment where females are likely to oviposit (Wang and Greenfield 1994; range = 9–14 host plants per site, median = 11). Additional details of photographic methods are in Supporting Information 1.

#### Image processing

We analyzed patterning in grasshopper body regions that would be visible in a resting pose. Specifically, we included the head, thorax, and femur, but excluded transparent wings because these transmitted the background color. We manually isolated stem images from their background and removed portions of the image that reflect damage from processing (e.g., bare wood at clipped ends of stem). Ground images were cropped to 15 cm^2^.

We used the Mica Toolbox v1.22 (Troscianko and Stevens 2015) in ImageJ (Abràmoff et al. 2004) to color-correct and rescale images, select regions of interest, and calculate image luminance under a model of human vision. While this measure of luminance does not include UV (which may be visible to avian predators) or model predator visual acuity (as in van den Berg et al. 2020), it provided a useful approximation of ecologically relevant luminance variation in grasshoppers and their environment.

#### Pattern quantification

We used fast Fourier transformation and bandpass filtering (Stoddard and Stevens 2010) to summarize patterning in grasshoppers and their environment. This approach quantifies the magnitude of luminance variation (“pattern energy”) that occurs over a range of discrete spatial scales, which can then be summarized in a pattern energy spectrum (Troscianko and Stevens 2015). Pattern energy at small spatial scales corresponds to fine-scale contrast (e.g., speckling), while higher scales correspond to larger contrasting features (e.g., blotches, heterogeneous gravel). We considered spatial scales from 0.12 —19.69 mm (bounded on the low end by camera sensor / lens combination and on the high end by grasshopper body size), with a multiplicative step size of 2^⅓^. We used the resulting pattern energy spectra to summarize pattern variation in substrate and stem samples using principal components analysis (PCA) in R v3.6.1 (R Core Team 2019)

#### Phenotype-environment correlations

We fit binomial generalized linear models (GLMs) to determine if geographic variation in morph frequencies was correlated with stem and/or ground patterning (summarized as mean PC scores for each site). We inferred the significance of model terms with likelihood-ratio χ^2^ tests with the Anova() function in the car package (Fox and Weisberg 2019) and estimated model *r*^2^ as in (Zhang 2017) with the rsq package (Zhang 2018) in R. For all models we considered the main effects of PCs that cumulatively accounted for ≥ 80% of pattern variation (PC1 and PC2 for substrate, PC1 for stems) and two-way interactions. We scaled PC scores (mean = 0, s.d. = 1) prior to analysis to facilitate interpretation.

### DNA sequencing and RAD assembly

#### Collections

We performed genetic analyses with 530 desert clickers collected across the species’ range from 2015—2018, of which 500 were scored for color morph phenotype (**Fig 2**). Of these, 298 individuals were used in analyses of phenotype-environment correlations and another 13 had photo vouchers. Color morphs for all individuals were subjectively scored by one of us (TKO) based upon vouchers and/or preserved specimens. In practice, banded morphs were recognized by a lighter band of cuticle at the margin of the pronotum, though the prominence of this band varied among individuals (see Results). All specimens were stored in 100% ethanol and stored at −20°C until DNA extraction. Further collection and DNA extraction details in Supporting Information 1.

**Figure 2.**
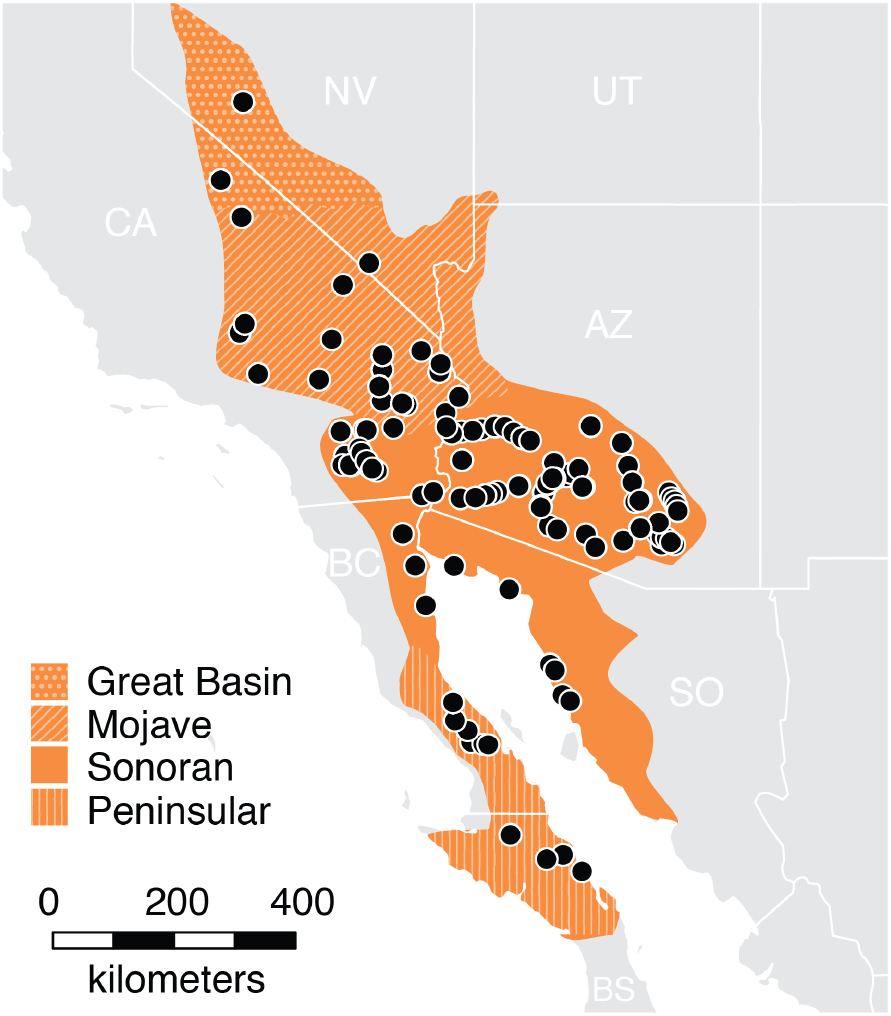
Localities of 530 collections made for this study (black dots) span most of the desert clicker’s distribution (orange). Pattern indicates desert region. CA: California. NV: Nevada. UT: Utah. AZ: Arizona. BC: Baja California. BS: Baja California Sur. SO: Sonora.

#### RADcap sequencing

We used reduced representation sequencing to begin to study the genetic basis of color polymorphism. RADcap (Hoffberg et al. 2016) sequencing pairs a double-digest RAD library preparation (3RAD, Bayona-Vásquez et al., 2019) with sequence capture for economical, repeatable, and high-throughput genotyping. This approach was attractive for our system given the size of grasshopper genomes (∼6–16 Gb, Gregory 2020) and the large number of samples in our dataset. We designed custom sequence capture baits targeting 10,097 loci identified from pilot double-digest RAD sequencing (Supporting Information 1). To reduce project cost, we combined desert clicker baits with 29,903 baits targeting other species and ordered them as a single pool from Arbor Biosciences (myBaits Custom kit). Although baits were at ∼0.25× concentration relative to manufacturer’s recommendations, RADseq libraries captured with 0.2× baits can perform as well as those captured with 1× baits (Ali et al. 2016)

We prepared 3RAD libraries as in Bayona-Vásquez et al. (2019) and performed sequence capture following manufacturer’s instructions, with minor adjustments (Supporting Information 1). We sequenced libraries across portions of two HiSeq 4000 lanes at the Vincent G. Coates Sequencing Laboratory at UC Berkeley.

#### RAD assembly

To generate artificial reference sequences for read mapping (“radnome” of Hoffberg et al. 2016), we found the consensus sequence of each target locus in our pilot double-digest RADseq (ddRAD) data. We then performed paired-end reference-based RAD assembly with ipyrad v.0.9.57 (Eaton and Overcast 2020). Assembly parameters were set to default except: restriction_overhang = TAATT, ATGCA; datatype = pair3rad; filter_adapters = 1; filter_min_trim_len = 80; max_SNPs_locus = 0.75; max_indels_locus = 10; max_shared_Hs_locus = 0.75. We output the assembly in .vcf format and used vcftools v0.1.15 (Danecek et al. 2011) to retain only biallelic SNPs with minor allele count ≥ 3 (Linck and Battey 2019).

The paired-end RAD assembly pipeline implemented in ipyrad required that R1 and R2 reads both map to a reference sequence. Cut-site polymorphisms that caused one read to go unmapped therefore resulted in missing data across an entire locus, even if the second read contained informative data. We reduced missingness by generating separate R1 and R2 assemblies, then, for each locus, retaining data from the more complete assembly. Sequence-based analyses used this optimized single-end assembly, while analyses of locus genotyping rates used both R1 and R2 assemblies to determine locus presence / absence in our dataset.

### Detecting structural variants with RADseq data

Recent insights into structural variation have been enabled by whole-genome resequencing and an evolving toolkit for structural variant detection (Alkan et al. 2011; Ho et al. 2020). While the resources to exhaustively characterize structural variants are becoming more attainable, most researchers of non-model species – especially those with large genomes – are still limited to reduced-representation sequencing approaches (e.g., variants of RADseq, Andrews et al. 2016). This technical and financial hurdle has limited the study of structural variants and their phenotypic consequences in non-model species.

We reasoned that large structural variants should leave distinctive signatures in RADseq data, allowing us to detect and characterize inversions and/or indels that differ between morphs even without a reference genome (**Fig 3**). We specifically considered the effect of inversions and indels on genotyping rates and genotype-phenotype associations.

**Figure 3.**
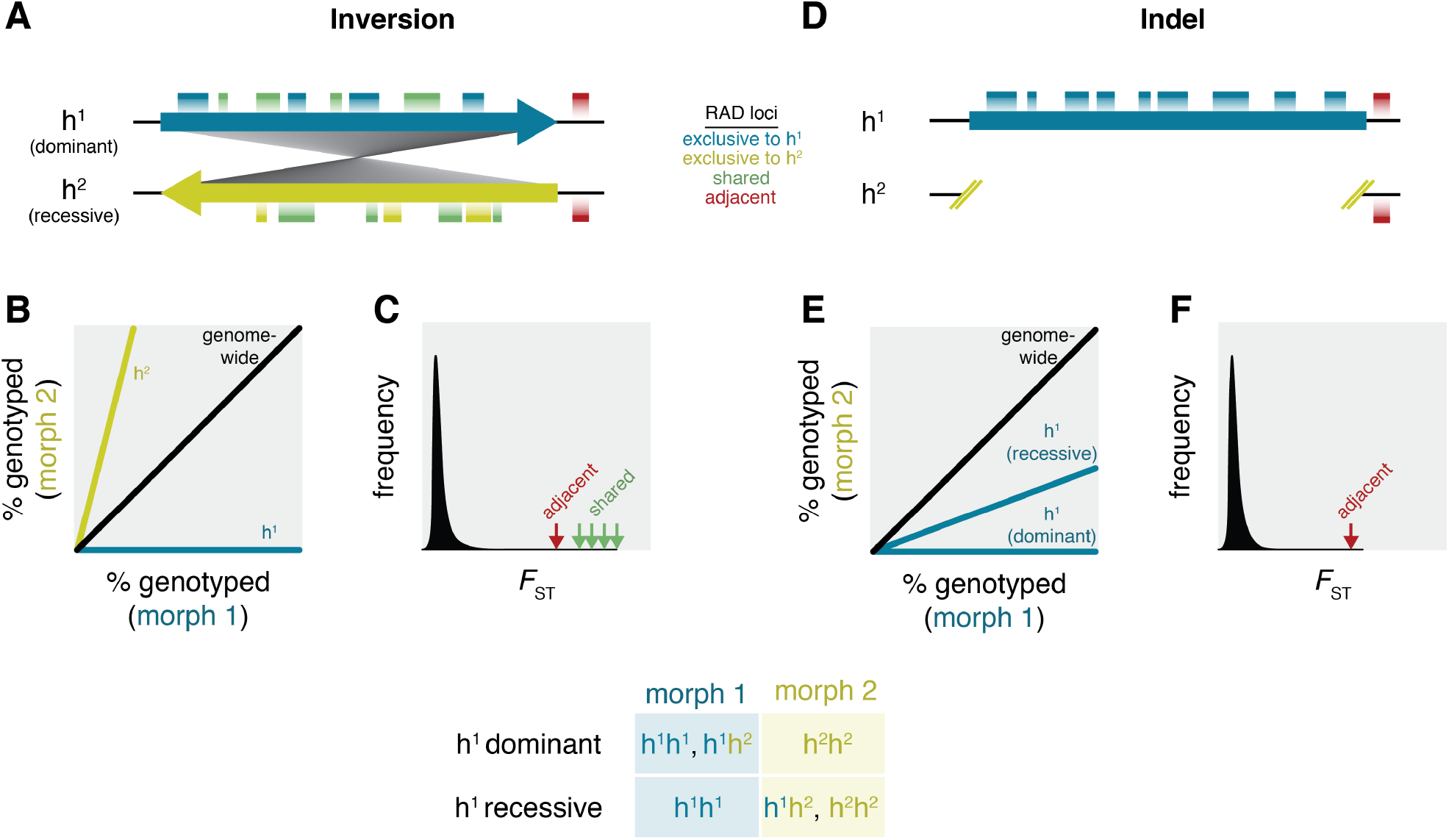
Predicted patterns of genotyping rates and between-morph *F*_ST_ if color morph is linked to a chromosomal inversion (A–C) or a large indel (D–F). Key at bottom shows possible genotypes for each morph if one haplotype is fully dominant. **A**. Cartoon of two inverted haplotypes, h^1^ and h^2^ (arrows). Due to common ancestry, some RAD loci are shared between haplotypes (green bars). However, divergence between inverted regions can eliminate shared restriction sites and create novel ones, resulting in RAD loci that are exclusive to each haplotype (blue and yellow bars). Suppressed recombination can also extend beyond inversion breakpoints to include adjacent RAD loci (red bars) **B**. Predicted differences genotyping rates between morphs. The vast majority of loci should be genotyped at comparable rates in each morph (black line). Loci exclusive to the dominant haplotype (h^1^) should be genotyped only in individuals the dominant phenotype (blue line), while loci exclusive to h^2^ should be genotyped at higher rates in morph 2 (yellow line). **C**. Hypothetical *F*_ST_ outlier scan. Shared loci within the inversion are expected to be exceptionally differentiated between morphs. Adjacent loci may also be divergent (red), but less so due to occasional recombination. **D**. Cartoon of two haplotypes that differ by a large indel. **E**. Loci exclusive to h^1^ will be genotyped only in morph 1 (if h^1^ is dominant) or predominantly in morph 1 (if h^1^ is recessive). **F**. As with inversions, loci adjacent to the indel may experience reduced recombination that elevates *F*_ST_ between morphs.

Assuming a collinear genome and random mating between morphs, each morph should be equally susceptible to the causes of missingness in RADseq datasets (e.g., restriction-site polymorphisms, insufficient sequencing coverage, or imperfect sequence capture design (Andrews et al. 2016; Hoffberg et al. 2016)). We therefore expect RAD loci to be genotyped at equal rates in each morph even if genotyping success is variable among loci. Mean *F*ST between morphs should be approximately zero across the genome, with the exception of loci linked to variants underlying color morph (Lewontin and Krakauer 1973). However, the power to detect genotype-phenotype associations with *F*ST outlier scans of sparsely sampled markers is low (Lowry et al. 2016).

Suppressed recombination within a chromosomal inversion (**Fig 3A**) allows each haplotype to fix alternative polymorphisms and accumulate novel mutations (Kirkpatrick 2010). Because haplotypes can gain or lose restriction sites independently, inverted regions should have complements of RAD loci that are exclusive to each haplotype as well as loci they share due to common ancestry. A subset of loci that are genotyped at drastically different rates in each morph can therefore reveal the presence of inversions (**Fig 3B**). Loci exclusive to the dominant haplotype will be found only in the dominant morph. Loci exclusive to the recessive haplotype will be genotyped in both morphs but overrepresented in the recessive morph. Inversion polymorphisms also have predictable effects on *F*ST outlier scans. RAD loci that are shared by both haplotypes should be exceptionally differentiated between morphs despite an otherwise homogeneous genomic background (Kirkpatrick 2010, **Fig 3C**). Because suppressed recombination extends beyond inversion breakpoints (Gong et al. 2005), RAD loci adjacent to inversions may also have elevated *F*ST.

By contrast, only one haplotype will have an exclusive set of RAD loci following an insertion or deletion (**Fig 3D**). Haplotype-exclusive loci should be genotyped in only one morph if the larger haplotype is dominant, or at greater rates in one morph if the larger haplotype is recessive (**Fig 3E**). As with inversions, large indels can suppress recombination over megabase scales (Morgan et al. 2017). RAD loci adjacent to an indel may thus be exceptionally differentiated between morphs (**Fig 3F**).

#### Delimiting focal populations

Population structure can confound tests of genotype-phenotype association (Price et al. 2006) and population genetic summary statistics (e.g., Hammer et al. 2003). We therefore used principal PCA to identify a set of focal populations with minimal population structure (median *F*ST = 0.013, max *F*ST = 0.041, estimated with the method of Weir and Cockerham 1984). Our tests for genotype-phenotype associations initially focused on this subsample of 236 phenotyped individuals (78 banded, 158 uniform) from the western Sonoran Desert (Supporting Information 1).

We then compared findings from the western Sonoran Desert to populations across the desert clicker’s distribution. We used discriminant analysis of principal components (DAPC, Jombart et al. 2010) to group samples into eight genetic clusters for further analysis (details in Supporting Information 1).

#### Comparing genotyping rates

For each RAD locus we separately calculated genotyping rates in banded and uniform grasshoppers. We then identified loci deviating from the expectation of equal genotyping success using Fisher’s exact tests with a 5% false discovery rate correction.

#### Identifying and characterizing morph-associated loci

We used an *F*ST outlier scan to identify RAD loci that were exceptionally differentiated between color morphs. We calculated weighted *FST* between banded and uniform individuals for all RAD loci genotyped in ≥ 15% of individuals from each (8,270 loci). We then calculated Tajima’s *D* for all loci. Positive *D* values in the upper tail of the genome-wide distribution are consistent with allele-frequency distortions due to balancing selection (Tajima 1989). All calculations were performed with vcftools v0.1.15. For loci of interest, we inferred statistical parsimony haplotype networks (Templeton et al. 1992) using the haploNet function in the pegas package (Paradis 2010) in R.

## RESULTS

### Color morphs are distinct but geographically variable

Consistent with our field experience and earlier observations (McNeill 1897; Rehn 1923), image analyses revealed two discrete color morphs in the desert clicker. Banded individuals were characterized by a peak of pattern energy at the ∼1 mm scale, while uniform individuals had weaker, flatter energy spectra (**Fig 4A**). However, the distinctness of color morphs also varied geographically (**Fig S2**.**1, S2**.**2)**. Banded and uniform individuals were least differentiated in populations where grasshoppers were darker overall (**Fig S2**.**3**). This suggests that melanism in some populations masks the lighter cuticle patches of banded individuals, reducing visual differences between morphs.

**Figure 4.**
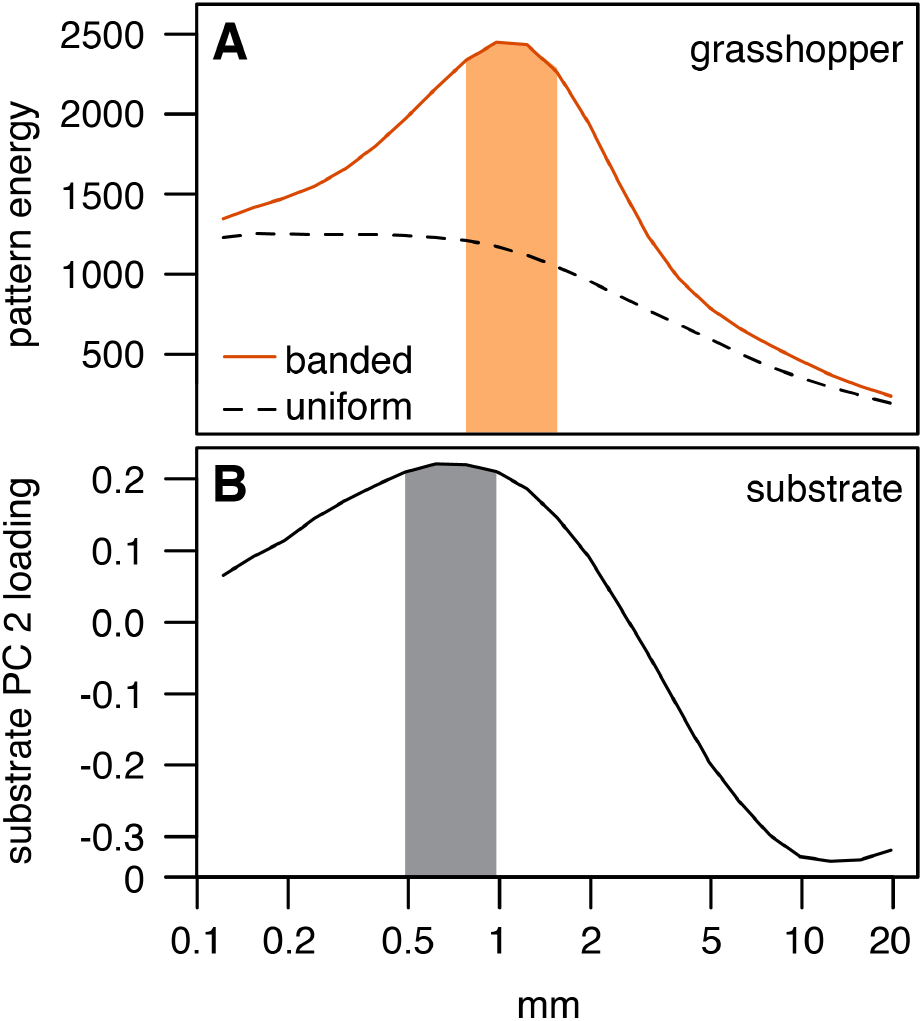
**A**. Comparison of mean pattern energy spectra for uniform (dashed black) and banded (solid orange) morphs. Banded morph patterning peaked at 0.75–1.25 mm, while uniform morphs had relatively flat energy spectra. **B**. Variable loadings on substrate PC2 peaked at 0.5–1 mm. Positive PC2 scores therefore indicated stronger patterning at this spatial scale. Area under the four highest values for each panel are highlighted to emphasize spectral peaks.

### Morphs coexist at intermediate frequencies

We identified both color morphs in 17 out of 19 populations. Banded individuals were typically found at intermediate frequencies, but were less common than the uniform morph (median = 29%, range = 10–79%; **Fig 5**). The two populations where we did not encounter banded grasshoppers had the fewest observations in our dataset (population C = 10, population M = 12), so it is possible that banded morphs occurred at low frequencies but were not encountered. Consistent with this hypothesis, we found banded individuals within 13 km of site C and across the full distribution of the desert clicker (**Fig S2**.**4**). The paucity of populations near fixation suggests that the balance of directional selection and homogenizing gene flow is unlikely to maintain color polymorphism.

**Figure 5.**
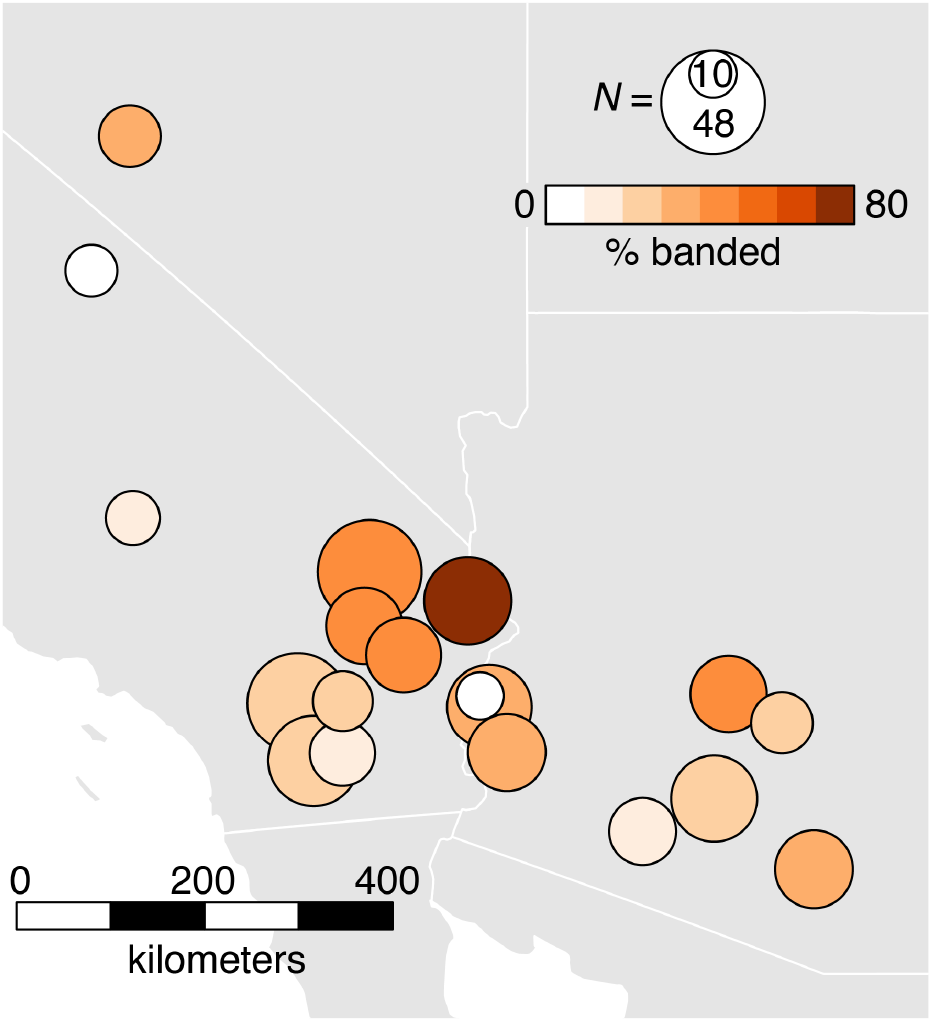
Banded morph frequencies in 19 focal populations.

### Desert substrate predicts local morph frequencies

Banded morph frequency covaried with visual features of desert substrate, but not host plant stems. A model including substrate PC1, substrate PC2, and their interaction explained the majority of variance in grasshopper morph frequencies (*r*^2^ = 0.60, **Table S2**.**1**). By contrast, stem PC1 did not predict morph frequencies alone (*r*^2^ = 0.02, **Table S2**.**2**) or through its interaction with substrate PCs (**Table S2**.**3**). The limited visual difference between banded and uniform morphs at three sites in eastern Arizona (sites R,S,T; **Fig S2**.**1**) suggests that differential selection on color morphs is probably weaker in these populations than elsewhere. When we excluded these three sites, we found an even stronger association between morph frequencies and substrate (*r*^2^ = 0.72, **Table 1**), but no correlation with stems (**Table S2**.**4**). Additional analyses focused on the model based with 16 sites.

**Table 1.** Summary of binomial generalized linear model of morph frequencies as a function of substrate patterning (mean PC scores). Significance of model terms was evaluated with a likelihood-ratio χ2 test with 1 d.f. Model *r*^2^ = 0.72. PC1 was a proxy for substrate contrast, while PC2 captured variation in the spatial scale of substrate pattern.

| | *B* | $\chi^2$ | *P* |
| --- | --- | --- | --- |
| substrate PC1 (contrast) | 0.73 | 35.8 | $2.2 \times 10^{-9}$ |
| substrate PC2 (scale) | -0.06 | 8.1 | $4.4 \times 10^{-3}$ |
| substrate PC1 × substrate PC2 | 0.46 | 11.1 | $8.8 \times 10^{-4}$ |

We examined variable loadings on substrate PCs to better understand the association between substrate pattern and morph frequencies. Substrate PC1 was a proxy for overall contrast (quantified as standard deviation of pixel luminance, *r*^2^ = 0.96). The strong positive effect of substrate PC1 on morph frequencies (*b* = 0.73) indicated that banded grasshoppers were more common in high-contrast substrates. The spatial frequencies that loaded positively on substrate PC2 (0.12–2.46 mm, peaking near 0.5–1 mm **Fig 4B**) closely corresponded to the spatial scale of light-dark patterning in banded grasshoppers (peaking near 0.75–1.25 mm, **Fig 4A**). While the main effect of substrate PC2 was weakly negative (*b* = −0.06), the strong and positive interaction with substrate PC1 (*b* = 0.46) revealed that substrate contrast best predicts morph frequencies when the substrate pattern resembles banded grasshoppers. That is, banded morphs are most common where they are likely to be most cryptic.

### A putative indel is associated with color morph

In the western Sonoran Desert, 18 out of 9,073 filtered RAD loci were genotyped at different rates in each color morph (**Fig S2**.**5**). Twelve of these were exclusively found in banded individuals (or nearly so), consistent with an indel polymorphism where the larger, dominant haplotype is absent from the recessive morph (**Fig 3E**). We refer to the larger haplotype as “B” and the smaller haplotype as “U” due to their association with banded and uniform morphs, respectively. Patterns of heterozygosity suggest that the putative indel occurs on an autosome. Only one sex may be heterozygous at a sex-linked locus under the sex determination systems typical of grasshoppers (XX/XO or neo-XY; Rodrigo et al. 2010). We instead found that a similar proportion of males and females were heterozygous at multiple indel loci (9/87 banded males, 6/23 banded females, Fisher’s exact test, *P* > 0.1).

Five additional loci were strongly overrepresented in banded morphs. Geographic signal in genotyping rates for four of these loci (**Fig S2**.**6**) suggests that the U haplotype is linked to cut-site polymorphisms in some populations, resulting in geographically-restricted null alleles (Andrews et al. 2016) and a bias towards genotyping banded individuals. A final locus was slightly overrepresented in uniform morphs, consistent with either linkage between the B haplotype and a cut-site polymorphism or a false positive. We did not identify a complement of loci that were found to both morphs but overrepresented in uniform individuals, as would be expected in the case of an inversion (**Fig 3B**).

We next imputed karyotypes for all individuals to determine whether the association between structural variation and color morph was consistent across populations. Karyotype assignments were based on genotyping success at the 12 indel loci. To guard against low levels of cross-contamination and index hopping (Illumina 2021), we required that individuals be genotyped at ≥ 2 indel loci to infer that they carried a B haplotype (karyotype BB or BU = B-). All other individuals were imputed as UU. Further details of inclusion criteria and imputation thresholds provided in **Fig S2**.**7**. After excluding 38 individuals with ambiguous or missing phenotypes and 50 individuals with ≥75% missing loci, we inferred putative karyotypes for a total of 442 individuals.

Karyotype-phenotype associations followed two distinct patterns across the desert clicker’s range (**Fig 6**). In the western Sonoran, northern Peninsular, Mojave, and Great Basin Deserts there was a near-perfect association between B-karyotypes and the banded phenotype (**Fig 6B, D, F**), indicating that B is dominant to U in these populations. The one exception was an individual from south-central Arizona (marked with a star in Fig 6A, B). All banded individuals from elsewhere in the range also had B-karyotypes (**Fig 6C, E, G**). However, 20% of uniform individuals had B-karyotypes in southern Baja California, Sonora, and eastern Arizona, indicating that many or all B haplotypes are recessive in these populations.

**Figure 6.**
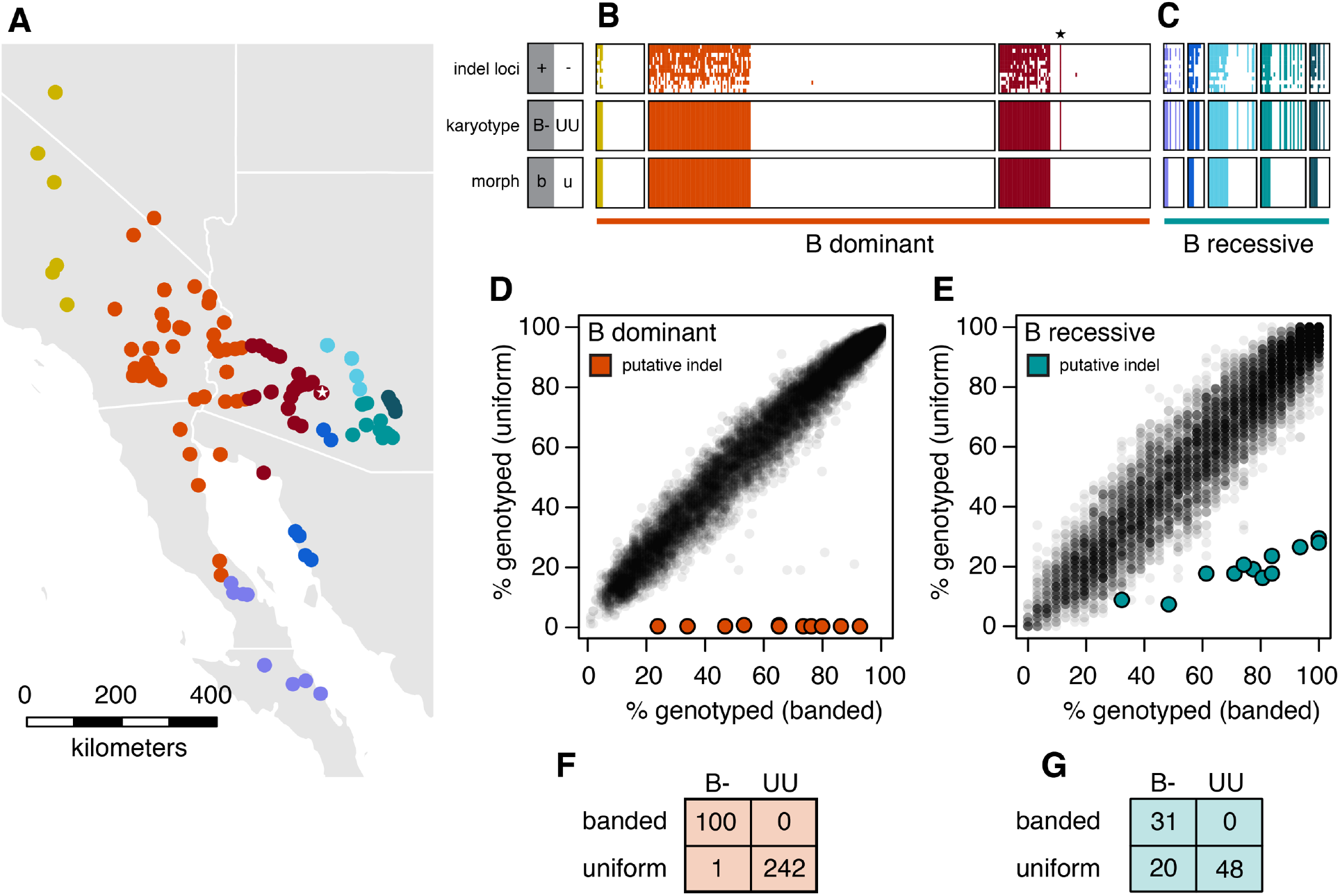
Dominance of putative indel differs among desert clicker populations. **A**. Distribution of eight population groups, coded by color. **B–C**. Correspondence of putative karyotype and phenotype across the desert clicker’s range. Populations grouped and colored as in panel A. Each column is a single individual. Top row: presence / absence of 12 indel loci in RAD dataset (color = present, white = absent). Middle: karyotype assignment based on presence / absence of 12 indel loci (color = B-, white = UU). Bottom: grasshopper morph (color = banded, white = uniform). **B**. The B-karyotype is nearly exclusive to banded individuals in the Great Basin, Mojave, western Sonoran, and northern Peninsular Deserts. The one exception (marked with a star) comes from the edge of the red population group in central Arizona (marked with a star in panel A). **C**. In the southern Peninsular and eastern Sonoran Desert, uniform morphs may be B- or UU. All banded morphs are B-. **D, E**. Between-morph differences in genotyping rates are consistent with a B haplotype that is dominant in some populations (D) and recessive in others I. Compare with Fig 2E. **F, G**. Tabular summary of data in panels B and C.

Dominance relationships did not differ between sexes across all populations (log-linear test for independence conditioned upon karyotype; χ^2^ = 0.67, d.f. = 2, *P* > 0.71) or when separately considering populations where B is recessive (χ^2^ = 2.01, 2 d.f., *P* > 0.35).

We next investigated the size and origin of the putative structural variant. First, we used patterns of between-locus linkage disequilibrium (LD) to determine if the 12 indel loci were linked (physically clustered) or freely recombining (found at the average genome-wide density). We calculated genotypic *r*^2^ between SNPs in indel loci with vcftools v1.15 (requiring minor allele frequency > 5% and < 80% missingness in B-individuals). We separately calculated *r*^2^ for two groups of populations in order to avoid the confounding effect of population structure (two-locus Wahlund effect). The first group included all western populations where B is dominant, while the second included three population clusters in eastern Arizona where B is recessive (see **Fig S1**.**5** for PCA of individuals labeled by genetic cluster).

Indel loci were unlinked in western populations where B is dominant. Median *r*^2^ between locus pairs ranged from 0.003 – 0.12 (median = 0.01, **Fig S2**.**8A**), indicating that indel loci are unlikely to be closely clustered on B haplotypes. If we assume the 12 indel loci are distributed at the expected genome-wide density (0.56–1.5 / Mb, for 9,000 loci across a typical grasshopper genome size of 6–16 Gb), we can roughly estimate the indel size to be 6.75–18 Mb. Between-locus LD was somewhat higher in eastern Arizona (*r*^2^ = 0.012 – 0.41, median = 0.10, **Fig S2**.**8B**), where population clusters are geographically isolated and may have lower population recombination rates due to reduced *N*e.

The putative structural variant may be the result of an insertion in the B haplotype or a deletion in the U haplotype. Horizontal gene transfer from mitochondria or bacteria (Wybouw et al. 2016) are potential sources of novel DNA, and *Wolbochia* in particular can occupy multiple Mb of insect chromosomes (Klasson et al. 2014). To test the hypothesis that indel DNA was horizontally transferred, we queried the consensus sequences from each of the 12 indel loci against the NCBI non-redundant (nr) database using DC-BLAST within the NCBI BLAST server (https://blast.ncbi.nlm.nih.gov/Blast.cgi). Search parameters were optimized for short queries (∼90 nt). We found no homology between the 12 markers and any sequence in the nr database (all E > 0.01), providing no support for horizontal gene transfer. In principle, localized proliferation of transposable elements may also generate a large insertions. We searched for repetitive elements in the 12 indel loci using the RepeatMasker web server (Smit, AFA, Hubley, R & Green, 2020) and Dfam 3.0 database (Hubley et al. 2016) but found none.

### Molecular signature of balancing selection on a morph-associated locus

We predicted that sites linked to the putative indel might also be associated with color morph due to extended LD near indel breakpoints. Consistent with this prediction, we identified a RAD locus that was strongly differentiated between morphs in the western Sonoran Desert (**Fig 7A**, weighted *F*ST = 0.56) in an otherwise homogeneous genetic background (mean weighted *F*ST = 0.002). A parsimony haplotype network inferred from 113 individuals revealed two major haplogroups at the morph-associated locus: one private to banded morphs and another shared by both morphs (**Fig 7B**).

**Figure 7.**
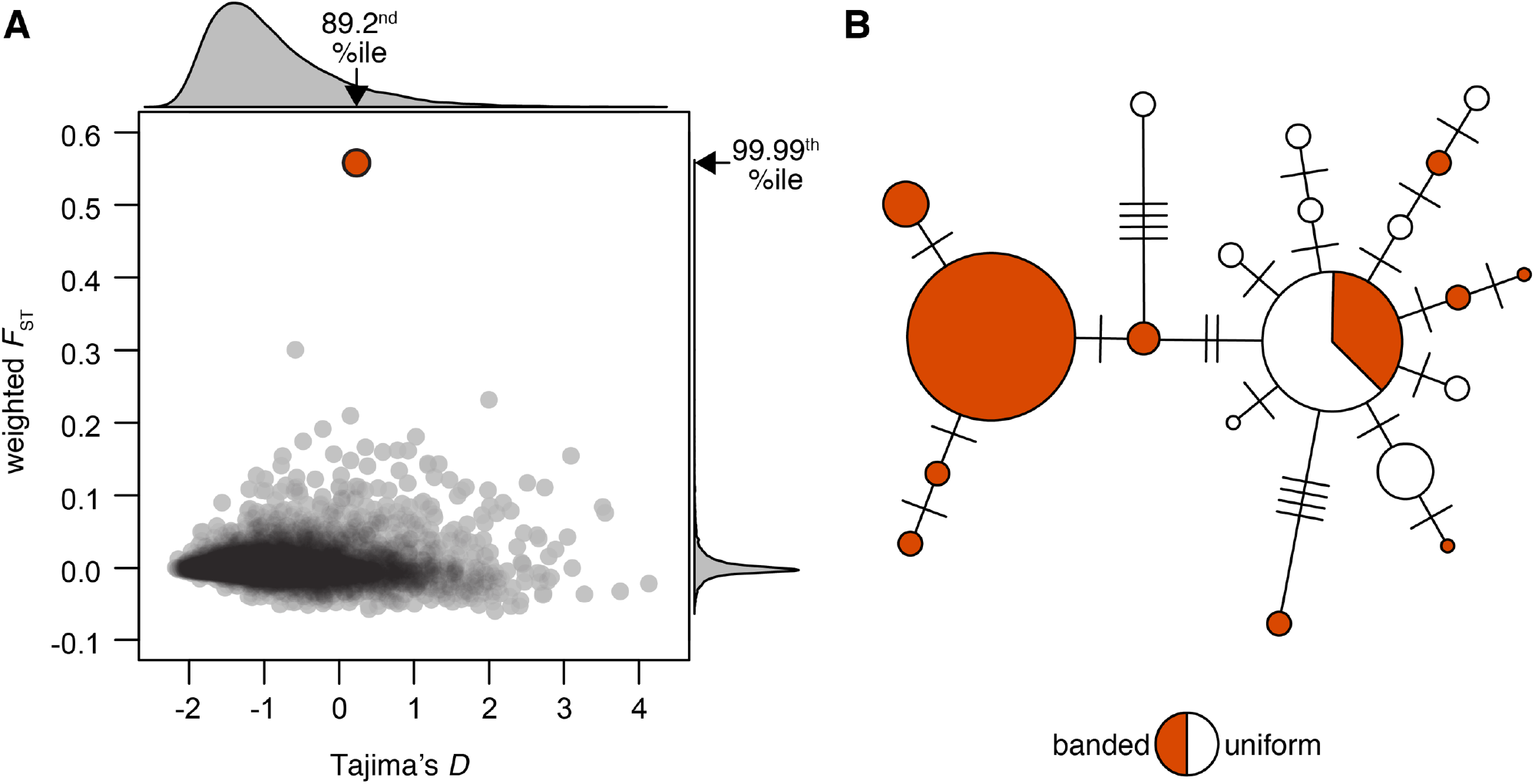
**A**. Genome-wide distribution of Tajima’s *D* and weighted *F*_ST_ between desert clicker morphs. One outlier (“morph-associated locus”) was strongly differentiated between morphs and found in the upper tail of the Tajima’s *D* distribution. **B**. Haplotype network of the morph-associated locus in the western Sonoran Desert. Circle size is proportional to number of haplotypes (max = 49, min = 1), and mutational steps are marked with hashes.

The association between phenotype and genotype at the morph-associated locus was imperfect. Nine of 41 banded individuals did not have an allele from the banded-only haplogroup and the association between this locus and color morph did not hold in other populations (**Fig S2**.**9**). Both observations are consistent with the effect of recombination that erodes linkage between the morph-associated locus and causative locus over time and space. Nevertheless, the morph-associated locus allowed us to perform a molecular test of the hypothesis that balancing selection maintains color morph variation. Tajima’s *D* for the morph-associated locus was positive and in the upper tail of the genome-wide distribution (*D* = 0.23, 89^th^ percentile, **Fig 7A**), suggesting that balancing selection at a linked site may have generated more even allele frequencies than expected under neutrality.

## DISCUSSION

Here, we examined the ecology and genetic basis of crypsis polymorphism in desert clickers by addressing two major hypotheses. First, we hypothesized that predator-mediated balancing selection maintains crypsis morphs within populations. Second, we hypothesized that crypsis variation is associated with structural variation, as has been recently described in many color-polymorphic species (Wellenreuther and Bernatchez 2018; Orteu and Jiggins 2020; Villoutreix et al. 2020). We found that color polymorphism is associated with a large putative indel and may be maintained by local balancing selection. Remarkably, the dominance of indel haplotypes differed among populations, a phenomenon for which there is a firm theoretical basis but few empirical examples. This work lays the foundation for future studies on the nature of balancing selection and mechanisms of dominance evolution in desert clickers. Furthermore, the approach we apply to identify putative structural variants may be widely applicable to other non-model taxa.

### Environmental correlates of morph frequencies

We were surprised to find that color morph frequencies were unrelated to variation in host plant stems. The absence of correlation between morph frequencies and stem patterning cannot be explained by the absence of predation, but may arise if stem predators do not differentiate between color morphs or if selection is consistent among sites. We instead found that desert substrate – an environment used almost exclusively by females – explains the majority of variation in morph frequencies. This suggests that predation on ovipositing females underlies geographic variation in desert clicker crypsis even though both sexes spend the majority of their lives in plants (Rehn 1923; Wang and Greenfield 1994). While anecdotal, it is noteworthy that a plasticine desert clicker model suffered an unambiguous rodent attack when left on the ground, but no such models were attacked on plant stems (**Fig S2**.**10**). Additional sources of unexplained variance may include geographic turnover in predator communities (e.g., Rönkä et al. 2020) or tradeoffs between crypsis and thermoregulation (e.g., Forsman 2018).

We cannot yet evaluate how each color morph confers crypsis in the environments where they are favored. One possibility is background matching (Endler 1978), whereby grasshoppers evade predation by resembling a sample of their environment. The color and speckling of uniform morphs are often remarkably similar to host plant stems (**Fig 1A**), and banded individuals may likewise resemble desert substrate or blotchy stems. Alternatively, the high-contrast patterns in banded morphs may create false body lines that impede predator detection in a complex visual environment (disruptive patterning, Stevens & Merilaita, 2009). The same color pattern may also provide different modes of crypsis in different contexts (Price et al. 2019). Distinguishing between alternative forms of crypsis is now possible with analysis of full-spectral images and models of predator vision (van den Berg et al. 2020).

### Balancing selection on color polymorphism

Color morphs coexist at intermediate frequencies across the desert clickers’ range and Tajima’s *D* was elevated at a morph-associated locus. Together, these findings suggest that local balancing selection maintains color polymorphism within populations. The correlation of morph frequencies and substrate patterning further indicates that the polymorphic equilibrium depends upon the local environment, as has been described in other systems (Ozgo 2011; Takahashi et al. 2011; McLean et al. 2015; Svensson et al. 2020). The specific form(s) of balancing selection that maintains crypsis polymorphism remain to be elucidated, however. Our results suggest several possibilities.

Many crypsis polymorphisms are thought to be maintained by apostatic selection, a form of predator-mediated negative frequency dependence (Cain and Sheppard 1954; Clarke 1962). Under this model, predators optimize foraging by preferentially searching for common prey morphs; the resulting fitness advantage of rarity maintains local polymorphism (reviewed in Bond 2007). Relative detectability against a given background affects the equilibrium frequency of each morph under apostatic selection (Bond and Kamil 1998), which predicts covariation between the visual environment and morph frequencies. Consistent with this model, banded morphs were most common where they more closely resembled the local substrate. Available data cannot address the role of predator behavior in polymorphism maintenance, however.

Alternatively, stem- and ground-foraging predators may select for different color morphs due to visual differences between their respective environments. Because only females are exposed to predation on the ground, this would amount to sexually antagonistic selection (Owen 1953; Haldane 1962; Bonduriansky and Chenoweth 2009; van Doorn 2009). A stable polymorphism under sexually antagonistic selection generally requires strong and opposing selection on each sex (Kidwell et al., 1977; Patten & Haig, 2009) – which is possible in this system – or sex-specific dominance (Gillespie 1978; Fry 2010; Jordan and Charlesworth 2012; Connallon and Chenoweth 2019) – which we did not find. Spatial variation in morph frequencies could arise if sex-specific selection pressure covaries with one or both of the predation environments. We cannot directly test this mechanism with our data, but its ecological basis is plausible. Sex-specific habitat use leads to opposing selection on color morphs in vipers, skinks, and pygmy grasshoppers (Forsman 1995;

Forsman and Shine 1995; Forsman and Appelqvist 1999) and may also account different morph frequencies in male and female Australian plague locusts (Dearn and Davies 1983). Our data are not consistent with Levene’s model of spatially varying selection (Levene 1953), which maintains color polymorphism in *Timema* stick insects (Sandoval 1994a,b). Levene showed that under some conditions, polymorphism can be maintained in a discretely patchy habitat when no morph is universally favored across patches. The relative size and selection coefficients of each patch set the local polymorphic equilibrium, such that variation in either factor can account for geographic variation in morph frequencies. This model would be plausible if morph frequencies covaried with stem patterning. Male desert clickers typically establish persistent territories on discrete host plants (Greenfield et al. 1989; Wang and Greenfield 1994) that differ in patterning, approximating Levene’s model. Instead, morph frequencies were associated with substrate patterning. To align with our findings, desert substrate must be discretely patchy at a fine spatial scale and females must remain within a single patch type for long periods. We find no support for these assumptions in previous work on desert clicker behavior (Wang and Greenfield 1994), our field experience, or by analyzing within-site substrate variation (**Fig S2**.**11**), Finally, it is possible that one or more mechanisms interact to jointly maintain polymorphism and determine local morph frequencies (Mokkonen et al. 2011; Chouteau et al. 2017). This may include additional forms of balancing selection such as overdominance or negative-assortative mating that cannot in themselves account for the observed phenotype-environment correlations.

### A large structural variant associated with color morph

Many recent studies have identified structural variants associated with animal color polymorphisms (Wellenreuther and Bernatchez 2018; Orteu and Jiggins 2020; Villoutreix et al. 2020), but because this work has been taxonomically restricted, it is unclear whether the link between color and structural polymorphisms holds across animal diversity. In light of the tremendous color and karyotypic variation found across grasshoppers, we hypothesized that a structural variant underlies color polymorphism in desert clickers. Methods for detecting structural variants have advanced alongside sequencing technologies in recent years (Alkan et al. 2011; Ho et al. 2020). However, these methods require reliable genome assemblies and resequencing data that were unavailable to us. We therefore defined an approach for detecting structural variants with data that are widely accessible in nearly any system (i.e., RADseq and related sequencing methods).

Our approach uses biased genotyping rates to identify structural variants that distinguish phenotypic classes, and can be applied to any species with a discrete polymorphism. The sensitivity of this method scales with marker density. For example, if three RAD loci with extreme genotyping bias are required to call a structural variant, then many published studies are powered to identify indels ≥ 750 kb in size (mean marker density of 1 per 245 kb in survey of (Lowry et al. 2016)). The ability to detect inversions will depend upon both the size and age of the variant, with old and large inversions being easiest to detect. We note that our approach cannot conclusively evince any particular structural variant, nor can it flag complex mutations such as translocations coupled with duplications, insertions, or deletions (Frederik et al. 2021). Nevertheless, it may be widely useful for initiating studies on the genetic basis of color polymorphism, alternative reproductive strategies, or life history polymorphism in non-model taxa with RADseq data.

Consistent with our hypothesis, we identified a putative indel associated with color morph in desert clickers. We estimate the indel accounts for ∼0.13% of the desert clicker genome (∼6.75–18 Mb) and may span ∼25 genes (assuming ∼19,000 total genes as in the desert locust *Schistocerca gregaria* (Verlinden et al. 2020)). The larger indel haplotype (B) was found in 100% of banded desert clickers (131/131), suggesting that the structural variant may include the loci that underlie color morph variation. This hypothesis remains to be tested.

Our results mirror recent findings in *Timema* stick insects, where green and brown morphs of Northern California *Timema* differ by a ∼ 5 Mb deletion (Villoutreix et al. 2020). The structural variant spans loci underlying continuous color variation in a sister lineage, demonstrating how large deletions can convert a quantitative character into a discrete one. Other indels linked to ecologically relevant polymorphisms include a 1.8 kb deletion in underlying pattern variation in *Heliconius cydno galanthus* (Gallant et al. 2014) and secondary deletion of a masculinizing supergene in *Oedothorax gibbosus* spiders (Frederik et al. 2021). The desert clicker now provides another likely case study.

Further work is required to confirm the genomic location of the putative indel, describe its structure, and infer its evolutionary origin. More granular investigations will be aided by a high-quality genome assembly and long-read sequencing (Ho et al. 2020), DNA-FISH (Osborne et al. 2001), and complementary cytogenetic approaches (White 1973). Comparisons with congeners *L. planum* from the Chihuahuan Desert and an undescribed species from Baja California (Otte 1981, TKO unpublished data) will help date the putative variant and polarize the indel. Notably, the undescribed *Ligurotettix* species from Baja California has a color polymorphism resembling that of the desert clicker (**Fig S2**.**12**). It will be interesting to determine whether the polymorphisms share a common genetic basis across species, and if so, whether long-term balancing selection or introgression can account for this trans-species polymorphism (Lindtke et al. 2017; Jamie and Meier 2020; Palmer and Kronforst 2020; Villoutreix et al. 2020).

### Dominance

Whether dominance evolves in response to natural selection is an enduring debate in population genetics. An early model posited that dominance at a primary locus can be altered by selection on modifier alleles at a secondary (epistatic) locus (Fisher 1928a,b). This view was refuted on both population genetic and biochemical grounds (Wright 1929, 1934; Haldane 1930), and arguments against it were elaborated in succeeding decades (reviewed in Porteous 1996; Mayo and Bürger 1997; Bourguet 1999; Bagheri 2006). A key objection was that selection on modifier alleles is too weak to overcome drift when the primary locus is at mutation-selection equilibrium (Wright 1929). On the other hand, modifier alleles can readily invade when heterozygotes are common (Wright 1929; Fisher 1930, 1931; Haldane 1956; Feldman and Karlin 1971; Charlesworth and Charlesworth 1975; Bürger 1983; Otto and Bourguet 1999). While the modifier model has been rejected as a general explanation for dominance relationships, it may bear on an important subset of loci under balancing selection or non-equilibrium conditions (Billiard and Castric 2011).

A handful of examples of dominance evolution in natural systems are consistent with the modifier model. As the frequency of melanic peppered moths increased during England’s industrialization, so did the dominance of the melanic *carbonaria* phenotype (Kettlewell 1955, 1961; Haldane 1956). Outcrossing diminished *carbonaria* dominance, suggestive of epistatic dominance modifiers in English populations (Kettlewell 1965). Loci underlying mimicry polymorphism in *Heliconius* (Le Poul et al. 2014) and *Papilio* (Clarke and Sheppard 1960; Nijhout 2003) butterflies often show complete dominance between alleles that naturally co-occur, but incomplete and/or mosaic dominance between alleles found in allopatry. This accords with theory that predicts selection will refine dominance relationships to maintain color pattern fidelity (Charlesworth and Charlesworth 1975; Llaurens et al. 2013). The evolution of pesticide resistance in *Culex* mosquitoes demonstrates dominance evolution in real-time: some variation in insecticide resistance cannot be explained by amino acid substitutions in target enzymes, implicating epistatic modifiers that alter enzyme expression levels (Bourguet et al. 1997). Recent work in plants (Brassicaceae) showed that *trans*-acting small RNAs underlie dominance variation at self-incompatibility loci (Tarutani et al. 2010; Yasuda et al. 2016), providing the first mechanistic evidence of epistatic dominance modifiers.

We found that dominance relationships between putative B and U haplotypes differed among desert clicker populations. The B haplotype conferred a dominant banded phenotype in much of the desert clickers’ range, where color morphs are strongly differentiated and populations show little genetic structure. By contrast, the banded phenotype was recessive and only weakly penetrant in a set of genetically and geographically isolated populations in Baja California, Sonora, and eastern Arizona. Based on this pattern, we speculate that the banded phenotype was ancestrally recessive but evolved dominance in part of the species’ range, possibly in concert with increased penetrance.

Although our results cannot identify the cause of dominance evolution, they are compatible with several testable hypotheses. First, selection may have acted on epistatic dominance modifiers as proposed by Fisher (Fisher 1928a,b). Second, selection may have acted directly upon a causative locus to favor the replacement of banded or uniform alleles with altered dominance (Haldane 1930). Allelic turnover – the replacement of one balanced allele by its descendent – has been shown for a number of systems (Takahata et al. 1992; Guillemaud et al. 1998; Lighten et al. 2017; Palmer and Kronforst 2020), though we are not aware of any case in which the replacement affects dominance. Third, the derived color morph may have evolved multiple times with differing dominance. However, it is improbable that a mutation spanning the same region would arise twice in independent populations. Finally, dominance of the banded phenotype may have evolved as a pleiotropic consequence of selection on the dominance of other phenotypes affected by the indel polymorphism (Gould and Lewontin 1979; Barrett and Hoekstra 2011). These hypotheses await further investigation.

Why altered dominance of either morph would be favored is not immediately clear. Theory of dominance evolution under spatially varying selection, overdominance, and Batesian mimicry have been developed (O’Donald and Barrett 1973; Otto and Bourguet 1999), but expectations differ based on the form of selection. Additional work is required to predict the outcome of other forms of balancing selection that may operate on desert clicker color polymorphism (e.g., negative frequency-dependent selection, sexually antagonistic selection) (Billiard and Castric 2011). Absent this theory, we speculate that the frequency of a dominant banded phenotype – which is typically the less common morph – could be more precisely regulated by predators if all banded alleles are visible to selection, keeping desert clicker populations nearer their fitness peak.

## CONCLUSION

The desert clicker provides a promising model for further dissecting processes that maintain genetic variation in natural populations, the genetic basis of adaptation, and the evolution of dominance. Our results accord with recent findings that structural variants are linked to color polymorphisms in many taxa and suggest that grasshoppers may be fruitful models for further exploring this link.

## Supporting information

S2. Supplementary Results

## DATA AVAILABILITY

Raw sequence data is deposited in NCBI under BioProject PRJNA718684. Additional data are deposited in Dryad (xxx).

## ACKNOWLEDGMENTS

We thank KM Yule, AE Baniaga, SM Lambert, UO García-Vásquez, NM Alexandre, LD O’Connor, CL Kivarkis, and KM O’Connor for field assistance. Collections in Mexico were made under SEMERNAT permit FAUT-0247, and we thank M Trujano-Ortega for assistance with permitting. Collections in the United States were made under permit (California Department of Fish and Wildlife permit #13751, California Department of Parks and Recreation #17-820-37, Bureau of Land Management permit #4180, and a right-of-way grant from the Arizona Land Department) or on public lands that did not require express permission. Parts of this research were conducted on University of California Natural Reserves, and we thank C Tracy (Boyd Deep Canyon Desert Research Center) and J André and T La Doux (Sweeney Granite Mountains Desert Research Center) for their assistance. We thank LL Smith and K Bi for advice on designing and implementing the RADcap protocol, and I Overcast for assistance with ipyrad. We thank MD Greenfield for discussion of *Ligurotettix* predators. Funding for this project was provided by the Society for the Study of Evolution Rosemary Grant Award, American Society of Naturalists Student Research Award, American Museum of Natural History Theodore Roosevelt Memorial Fund, Orthopterists’ Society Theodore Cohn Research Award, Southwest Association of Naturalists Howard McCarley Research Award, UC Berkeley Essig Museum Walker Fund, UC Department of Integrative Biology, Philomathia Foundation Graduate Fellowship, and National Science Foundation Graduate Research Fellowship to TKO, as well as a grant from and National Institute of General Medical Sciences of the National Institutes of Health to NKW (R35GM119816).

## Notes

**Conflict of Interest Statement:** The authors declare no conflict of interest.

### Competing Interest Statement

The authors have declared no competing interest.

