## Supplementary material for "Ecological basis and genetic architecture of crypsis polymorphism in the desert clicker grasshopper (*Ligurotettix coquilletti*)": S1. Supplementary Methods

### SUPPORTING INFORMATION 1: Supplementary Methods

#### Phenotype and environmental survey

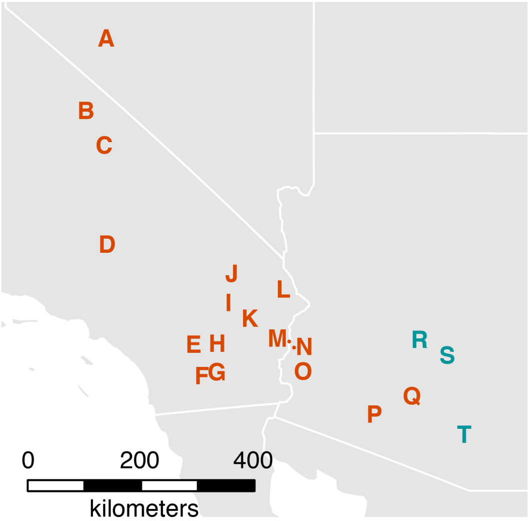


**Figure S1.1.** Location of 20 sites where we surveyed grasshopper phenotype and patterning of the visual environment. Site codes are centered over sampling site with the exception of M and N (points next to letters are sample locations). Orange = B haplotype inferred to be dominant, blue-green = B haplotype inferred to be recessive (see Results in main text). Site C was excluded from analysis due to the low number of desert clicker observations ($N$ = 4). At most sites (C-T) creosote bush (*Larrea tridentata*) was the primary host plant. Greasewood (*Sarcobatus vermiculata*) was the host plant at site A, and saltbrush (*Atriplex* spp.) was the host plant at site B.

**Table S1.1.** Coordinates of 20 sites represented in Fig S1.1.

| Site | Description | State | County | Latitude | Longitude |
|---|---|---|---|---|---|
| A | Mina | NV | Mineral | 38.5131 | -118.0737 |
| B | Bishop | CA | Inyo | 37.3609 | -118.4635 |
| C | Independence | CA | Inyo | 36.8139 | -118.1008 |
| D | Cinco | CA | Kern | 35.2423 | -118.0433 |
| E | Boyd Deep Canyon Desert Res Ctr | CA | Riverside | 33.6545 | -116.3743 |
| F | Ocotillo Wells | CA | San Diego | 33.1582 | -116.2113 |
| G | Salton City | CA | Imperial | 33.2208 | -115.9203 |
| H | Cactus City | CA | Riverside | 33.6734 | -115.9164 |
| I | Amboy Rd | CA | San Bernardino | 34.3189 | -115.6984 |
| J | Sweeney Granite Mountains Desert Res Ctr | CA | San Bernardino | 34.7807 | -115.6442 |
| K | Twentynine Palms Hwy near CA 177 | CA | San Bernardino | 34.0681 | -115.2998 |
| L | Turtle Mountain Rd | CA | San Bernardino | 34.5331 | -114.6502 |
| M | North of Blythe | CA | Riverside | 33.7186 | -114.5257 |
| N | Tom Wells Rd | AZ | La Paz | 33.6224 | -114.4316 |
| O | Kofa | AZ | Yuma | 33.2295 | -114.2543 |
| P | Goldwater Range | AZ | Maricopa | 32.5507 | -112.8790 |
| Q | Tabletop | AZ | Pinal | 32.8317 | -112.1542 |
| R | Rock Tank | AZ | Maricopa | 33.7348 | -112.0081 |
| S | Youngberg | AZ | Pinal | 33.4843 | -111.4676 |
| T | Tucson Mountains | AZ | Pima | 32.2280 | -111.1430 |

**Photography**

We captured all photographs in RAW format with a Nikon D7500 camera mounted 60 cm above each subject. To allow subsequent size and color calibration, we began each imaging session by photographing an XRite ColorChecker and scale bar under the same lighting conditions used for illuminating subjects.

We used a lightbox fashioned from a 40 $cm^3$ mesh cage and off-camera flashes to evenly illuminate images. Two opposing vertical faces of the lightbox were covered with polyester batting, which transmitted and diffused light from two Yongnou 560 Speedlites. Lights were fitted with gel diffusers and oriented at 15˚ above horizontal. The other vertical faces were covered in reflective white foamboard. The top face of the lightbox was also covered with foamboard, with a 10 cm x 15 cm hole to permit overhead photography. The bottom face was open so the lightbox could be placed over subjects.

*Substrate*

We photographed a ~20 cm x 30 cm patch of desert substrate 1 m from the base of ≥10 host plants per site (total $N = 222$). We minimized the effect of shadows caused by ambient sunlight by casting a shadow over the entire lightbox, so that artificial light dominated and natural light was diffusely distributed across the substrate. Photos were captured with a 40mm Nikkor Micro f/2.8 lens at f/22 (to maximize depth of field).

*Stems*

We photographed a total of 1004 stems in the laboratory against a low-reflection black velvet background. All stems were stored under cool conditions for up to two days until photographed. Photos were captured with a 40mm Nikkor Micro f/2.8 lens at f/10.

*Grasshoppers*

We maintained live grasshoppers in captivity until imaging (1-3 days). We immobilized grasshoppers by briefly chilling them in an ice-packed cooler or -20˚C freezer immediately before imaging. All grasshoppers were photographed in lateral view against a black velvet background with a Tamron 90mm f/2.8 lens at f/10.

### RADcap sequencing

#### *DNA extraction*

We extracted DNA from all samples using single-tube or 96-well DNeasy Blood and Tissue Kits (Qiagen). For adults and late-instar juveniles, we rehydrated one femur in autoclaved ddH20, chilled it to -80˚C, then homogenized in a bead mill for 1-4 minutes at 25-50 Hz. For early instar juveniles we homogenized whole bodies. We followed manufacturer's protocols with the following exceptions: samples were digested with proteinase K overnight and treated with RNAse A prior to binding to silica columns.

#### *Pilot ddRAD sequencing*

We performed pilot double-digest RAD (ddRAD) sequencing to determine whether standard reduced-representation protocols were suitable for genotyping hundreds of desert clickers. We prepared sequencing libraries for 27 desert clickers from Arizona, Nevada, and California (including both banded and uniform individuals) as in Peterson et al., (2012) with slight modifications to adapter design (see Barker et al., 2017). Grasshopper genomes are typically large (~6-16 Gb, Gregory 2020) and repetitive (Wang et al. 2014; Verlinden et al. 2020), so digestion with frequent-cutting enzymes would result in an excess of sequenced loci and reduced coverage for a given sequencing effort (Peterson et al. 2012). We attempted to limit the number of RAD loci by using two 6-cutter enzymes (EcoRI and PstI) and selecting a narrow size range of digested fragments for sequencing with a Pippin Prep (375-475 bp). We then sequenced each library on 1/96 of an Illumina HiSeq 4000 lane (PE 100) and performed QC and de-novo RAD assembly with ipyrad v0.7.19 (Eaton 2014). Loci were clustered at 90% identity, and we retained loci found in ≥ 2 individuals. We used default values for all other assembly parameters.

Despite efforts to minimize the number of sequenced regions, we assembled ~385,000 RAD loci with low locus coverage across samples. The high degree of missing data indicated that the sequencing effort required for a large-scale population genomics study with ddRAD would be cost-prohibitive. However, these pilot data allowed us to design custom sequence capture baits that would enable a larger scale sequencing effort.

#### *Bait design*

In order to economically genotype hundreds of samples, we performed RADcap sequencing (Hoffberg et al., 2016). RADcap combines 3RAD – a variant of double-digest RAD sequencing (Bayona-Vásquez et al. 2019) – with targeted sequence capture to enable reliable, efficient, and high-throughput sequencing of RAD loci. This approach presented several advantages for our project. First, combinatorial indexing (two in-line barcodes and one index read) permits massive multiplexing with minimal startup cost. Second, an 8 bp unique molecular index facilitates the identification and removal of PCR duplicates. Third, sequence capture increases the repeatability of genotyping across individuals and minimizes the sequencing required for sufficient coverage (Ali et al. 2016; Hoffberg et al. 2016).

We designed sequence capture baits from our pilot RAD assembly using a custom pipeline coded in Perl. We began by identifying suitable targets from among the full set of RAD loci assembled for each species, retaining loci that met the following criteria:

1) The locus was sequenced in at least two samples.

2) The locus lacked an NsiI recognition site, as this restriction enzyme was used to cleave adapter dimers during 3RAD library preparation (see below). Loci with the NsiI recognition site would be digested during library preparation and unsequenceable.
3) No sequenced alleles were > 5% divergent from the consensus sequence of that locus, as capture efficiency declines precipitously above 5% divergence (Bi et al. 2012).
4) The locus contained no indels, which can impede bait binding (Arbor Biosciences, 2017).
5) The locus consensus sequence had GC content between 30% and 70%, as extreme GC content can result in poor sequence capture efficiency (Bi et al. 2012).

We then performed an all vs. all BLAST search of retained loci and removed any locus with a significant hit using a liberal significance threshold (e-value ≤ 10). We further removed loci containing TE or simple repeat motifs identified with RepeatMasker v4.0.7 (Smit et al., 2015) and the RepBase arthropod repeat library (Bao et al. 2015). These steps minimized the possibility that baits would cross-bind or capture non-target loci with similar repeat motifs.

As in Hoffberg et al. (2016), we intended to combine bait sets for multiple species into a single custom bait order to minimize project cost. We used the same methods to identify target loci for three additional species from separate projects. Next, we performed an additional all vs. all BLAST search of loci from all four species and removed those with significant hits. Finally, we randomly selected 10,097 loci from the desert clicker and ~10,000 loci for three other species, then designed a single 80-bp bait sequence that was complementary to the consensus sequence for that locus.

We ordered a set of 40,000 myBaits sequence capture baits from Arbor Biosciences (Ann Arbor, Michigan), which included ~10,000 baits for each of four species. Baits for desert clicker loci were therefore at 0.25× concentration relative to the manufacturer's recommendation. Using an approach similar to RADcap (Rapture), Ali et al. (2016) found that capturing RAD loci with 0.2× bait concentrations can yield sequencing performance that is indistinguishable from libraries captured with 1× baits.

***Library preparation***
We designed custom adapters to match our set of restriction enzymes (PstI and EcoRI) and prepared 3RAD sequencing libraries following guidelines in (Bayona-Vásquez et al. 2019) (Table S3.1, Table S3.2). Prior to digestion, we quantified DNA concentration of each sample with Quant-iT PicoGreen dsDNA kit (Invitrogen) and a BioTek Synergy microplate reader. After diluting all samples to 10 ng / µL, we set up an initial digestion for all samples. Digestions consisted of: 10 µL DNA, 0.5 µL PstI-HF (NEB), 0.5 µL EcoRI-HF (NEB), 0.5 µL NsiI-HF (NEB), and 1.5 µL 10× CutSmart buffer (NEB). We then added a unique pair of barcoded adaptors to each sample defined its position in the 96-well plate. Eight unique "NsiI" adaptors (which ligate to DNA cut by PstI but are cleaved by NsiI if self-ligated) corresponded to the plate's eight rows, while twelve "EcoRI" adapters (which ligate to DNA cut by EcoRI) corresponded to the twelve columns (as detailed in Bayona-Vásquez et al., 2019). We digested DNA at 37˚C for 1 hour, then added the following ligation mixture to each sample: 2.75 µL molecular grade water, 1.5 µL ATP (NEB), 0.5 µL 10× ligase buffer (NEB), and 0.25 µL DNA ligase 400 u/mL (NEB). The combined digestion / ligation mixture was then cycled twice between ligation and digestion conditions (22˚C for 20 min, 37˚C for 10 min, 22˚C for 20 min, 37˚C for 10 min) before heat-killing the enzymes for 20 min at 80˚C.

We retained 10 µL from each reaction as a backup, then pooled the remaining 10 µL from all samples within a plate. Each plate was pooled separately. We cleaned pooled reactions with 1.25× SeraMag beads (prepared as in Rohland and Reich 2012) and resuspended samples in a total volume of 60µL molecular grade water.

For each pool, we next performed 6 replicate single-cycle PCR with an "iTru5 8N" primer containing a random 8 bp sequence and partial Illumina P5 adaptor (see Bayona-Vásquez et al., 2019). Each reaction contained: 25 µL KAPA HiFi HotStart ReadyMix (Roche), 10 µL pooled and cleaned DNA, and 5 µL iTru5 8N primer at 5µM. The single reaction cycle was as follows: 98˚C for 1 min, 60˚C for 30 sec, 72˚C for 6 min. We pooled replicate reactions and cleaned with 1.5× SeraMag beads, resuspending in 33 µL molecular grade water.

We next completed the Illumina adaptors and amplified libraries with triplicate reactions for each pool of DNA. Reactions consisted of: 25 µL KAPA HiFi HotStart ReadyMix (Roche), 5.0 µL P5 primer, 5.0 µL "iTru7" primer with an 8 bp plate-level barcode (see Bayona-Vásquez et al., 2019 for primer sequences), and 10 µL DNA resulting from pooled and cleaned single-cycle PCRs. Reaction conditions were: 98˚C for 2 min, followed by 12 cycles of 98˚C for 20 sec, 60˚C for 15 sec, 72˚C for 30 sec, and a final 72˚C elongation for 5 min. We pooled and cleaned reactions with 1.5× SeraMag beads, resuspending in 60 µL.

We quantified resulting DNA libraries with the Qubit HS dsDNA kit (Invitrogen) and calibrated the quantity of DNA to use in sequence capture. We expected the ratio of target to non-target DNA in our samples to be low, since our baits targeted a minority of RAD loci generated by double-digestion. In addition, the reduced relative concentration of baits further limited the opportunities for bait binding. In such conditions, increasing total amount of DNA in capture reactions can improve capture performance (McCartney-Melstad et al. 2016, Arbor Biosciences 2018). We therefore used between 1.5 and 2 µg of DNA per capture reaction.

We performed sequence capture following manufacturer's instructions (myBaits protocol v4, Arbor Biosciences) with two deviations. First, we used 10 µL Roche SeqCap EZ Developer Reagent (Roche) in lieu of Block C and Block O provided with the myBaits kit. For invertebrates, the developer reagent may provide better blocking of repetitive DNA (Ke Bi, personal communication). DNA mixed with 10 µL Developer Reagent and 0.5 µL Block A (provided with the myBaits kit) were dried to a volume of 7.5 µL in a SpeedVac prior to mixing with the hybridization mixture (step 1.4 in myBaits protocol v4). Second, we hybridized baits to DNA for 52 hours at 65˚C.

We directly amplified bead-bound libraries following capture. Triplicate reactions consisted of 25 µL KAPA HiFi HotStart ReadyMix (Roche), 5.0 µL P5 primer, 5.0 µL "iTru7" primer with 8 bp plate-level barcode, and 10 µL DNA. Reaction conditions were: 98˚C for 2 min, followed by 16 cycles of 98˚C for 20 sec, 60˚C for 15 sec, 72˚C for 30 sec, and 72˚C elongation for 5 min.

All samples for this project and several additional projects were sequenced across portions of two Illumina HiSeq 4000 lanes (PE100) at the Vincent J. Coates Sequencing Laboratory at the University of California, Berkeley. In total, we pooled 480 or 609 individuals per lane at equimolar concentrations.

**Table S1.1.** Oligo sequences for creating barcoded i5 adapter stubs following the protocols of (Bayona-Vásquez et al. 2019). "*" indicates phosphorothioate bond to prevent exonuclease degradation. "/5phos/" indicates 5' phosphorylation. These adapters ligated to desert clicker DNA digested by PstI, and adapter dimers were cleaved by NsiI.

| | | |
|---|---|---|
| i5-upper | iTru_NsiI_R1_stub_A | ACGACGCTCTTCCGATCTCCGAATATGC*A |
| i5-upper | iTru_NsiI_R1_stub_B | ACGACGCTCTTCCGATCTTTAGGCAATGC*A |
| i5-upper | iTru_NsiI_R1_stub_C | ACGACGCTCTTCCGATCTAACTCGTCATGC*A |
| i5-upper | iTru_NsiI_R1_stub_D | ACGACGCTCTTCCGATCTGGTCTACGTATGC*A |
| i5-upper | iTru_NsiI_R1_stub_E | ACGACGCTCTTCCGATCTGATACCATGC*A |
| i5-upper | iTru_NsiI_R1_stub_F | ACGACGCTCTTCCGATCTAGCGTTGATGC*A |
| i5-upper | iTru_NsiI_R1_stub_G | ACGACGCTCTTCCGATCTCTGCAACTATGC*A |
| i5-upper | iTru_NsiI_R1_stub_H | ACGACGCTCTTCCGATCTTCATGGTCAATGC*A |
| i5-lower | iTru_NsiI_R1_RCp_A | /5phos/T*ATTCGGAGATCGGAAGAGCGTCGTGTAGGGAAAGAGTGT |
| i5-lower | iTru_NsiI_R1_RCp_B | /5phos/T*TGCCTAAAGATCGGAAGAGCGTCGTGTAGGGAAAGAGTGT |
| i5-lower | iTru_NsiI_R1_RCp_C | /5phos/T*GACGAGTTAGATCGGAAGAGCGTCGTGTAGGGAAAGAGTGT |
| i5-lower | iTru_NsiI_R1_RCp_D | /5phos/T*ACGTAGACCAGATCGGAAGAGCGTCGTGTAGGGAAAGAGTGT |
| i5-lower | iTru_NsiI_R1_RCp_E | /5phos/T*GGTATCAGATCGGAAGAGCGTCGTGTAGGGAAAGAGTGT |
| i5-lower | iTru_NsiI_R1_RCp_F | /5phos/T*CAACGCTAGATCGGAAGAGCGTCGTGTAGGGAAAGAGTGT |
| i5-lower | iTru_NsiI_R1_RCp_G | /5phos/T*AGTTGCAGAGATCGGAAGAGCGTCGTGTAGGGAAAGAGTGT |
| i5-lower | iTru_NsiI_R1_RCp_H | /5phos/T*TGACCATGAAGATCGGAAGAGCGTCGTGTAGGGAAAGAGTGT |

**Table S1.2.** Oligo sequences for creating barcoded i7 adapter stubs following the protocols of (Bayona-Vásquez et al. 2019). "*" indicates phosphorothioate bond to limit nuclease degradation. "/5phos/" indicates 5' phosphorylation. These adapters ligated to desert clicker DNA digested by EcoRI.

| | | |
|---|---|---|
| i7-upper | iTru_EcoRI_R2_RC_stub_1 | A*ATTACGTTAGAGATCGGAAGAGCACACGTaatcc |
| i7-upper | iTru_EcoRI_R2_RC_stub_2 | A*ATTAGTACCGAAGATCGGAAGAGCACACGTaatcc |
| i7-upper | iTru_EcoRI_R2_RC_stub_3 | A*ATTACAACGATCAGATCGGAAGAGCACACGTaatcc |
| i7-upper | iTru_EcoRI_R2_RC_stub_4 | A*ATTAAGTGTAGCTAGATCGGAAGAGCACACGTaatcc |
| i7-upper | iTru_EcoRI_R2_RC_stub_5 | A*ATTAATGCGTAGATCGGAAGAGCACACGTaatcc |
| i7-upper | iTru_EcoRI_R2_RC_stub_6 | A*ATTATGCATACAGATCGGAAGAGCACACGTaatcc |
| i7-upper | iTru_EcoRI_R2_RC_stub_7 | A*ATTAGACATGTGAGATCGGAAGAGCACACGTaatcc |
| i7-upper | iTru_EcoRI_R2_RC_stub_8 | A*ATTATCGTGCACAAGATCGGAAGAGCACACGTaatcc |
| i7-upper | iTru_EcoRI_R2_RC_stub_9 | A*ATTATGATGCAGATCGGAAGAGCACACGTaatcc |
| i7-upper | iTru_EcoRI_R2_RC_stub_10 | A*ATTAACAGCATAGATCGGAAGAGCACACGTaatcc |
| i7-upper | iTru_EcoRI_R2_RC_stub_11 | A*ATTAAGGTCATGAGATCGGAAGAGCACACGTaatcc |
| i7-upper | iTru_EcoRI_R2_RC_stub_12 | A*ATTACTCACTGCAAGATCGGAAGAGCACACGTaatcc |
| i7-lower | iTru_EcoRI_R2_1 | GTGACTGGAGTTCAGACGTGTGCTCTTCCGATCTCTAACG*T |
| i7-lower | iTru_EcoRI_R2_2 | GTGACTGGAGTTCAGACGTGTGCTCTTCCGATCTTCGGTAC*T |
| i7-lower | iTru_EcoRI_R2_3 | GTGACTGGAGTTCAGACGTGTGCTCTTCCGATCTGATCGTTG*T |
| i7-lower | iTru_EcoRI_R2_4 | GTGACTGGAGTTCAGACGTGTGCTCTTCCGATCTAGCTACACT*T |
| i7-lower | iTru_EcoRI_R2_5 | GTGACTGGAGTTCAGACGTGTGCTCTTCCGATCTACGCAT*T |
| i7-lower | iTru_EcoRI_R2_6 | GTGACTGGAGTTCAGACGTGTGCTCTTCCGATCTGTATGCA*T |
| i7-lower | iTru_EcoRI_R2_7 | GTGACTGGAGTTCAGACGTGTGCTCTTCCGATCTCACATGTC*T |
| i7-lower | iTru_EcoRI_R2_8 | GTGACTGGAGTTCAGACGTGTGCTCTTCCGATCTTGTGCACGA*T |
| i7-lower | iTru_EcoRI_R2_9 | GTGACTGGAGTTCAGACGTGTGCTCTTCCGATCTGCATCA*T |
| i7-lower | iTru_EcoRI_R2_10 | GTGACTGGAGTTCAGACGTGTGCTCTTCCGATCTATGCTGT*T |
| i7-lower | iTru_EcoRI_R2_11 | GTGACTGGAGTTCAGACGTGTGCTCTTCCGATCTCATGACCT*T |
| i7-lower | iTru_EcoRI_R2_12 | GTGACTGGAGTTCAGACGTGTGCTCTTCCGATCTTGCAGTGAG*T |

**Identifying focal populations for genotype-phenotype associations**
To limit the confounding effect of population structure on tests of genotype-phenotype association, we attempted to identify a contiguous set of desert clicker populations with minimal genetic differentiation. We began by randomly selecting a single SNP per RAD locus that passed filtering (9,073 unlinked SNPs) and iteratively removing samples and loci with low coverage. The final dataset required all samples to be genotyped ≥ 70% of SNPs, and all SNPs to be genotyped in ≥ 95% of samples (482 individuals, 1,581 SNPs). Missing data for each locus were imputed with mean allele frequency. We then performed a principal components analysis on allele frequencies to summarize population structure.

Population structure in the western Sonoran Desert appeared to be relatively continuous, with limited isolation by distance (**Fig S1.2**). To further characterize these populations, we identified a cluster of 236 phenotyped individuals (78 banded, 158 uniform) and calculated pairwise $F_{ST}$ (Weir and Cockerham 1984)among 14 populations with four or more individuals (range = 4-21, median = 11.5). $F_{ST}$ was low among populations (median = 0.013, max = 0.041) consistent with minimal population differentiation. We therefore used these 236 individuals to initially test for genotype-phenotype associations, then compared these findings to other populations.

Analyses were performed using functions in the R packages vcfR v1.12.0 (Knaus and Grünwald 2016), adegenet v2.1.3 (Jombart 2008), hierfstat v0.5-7 (Goudet and Jombart 2020)

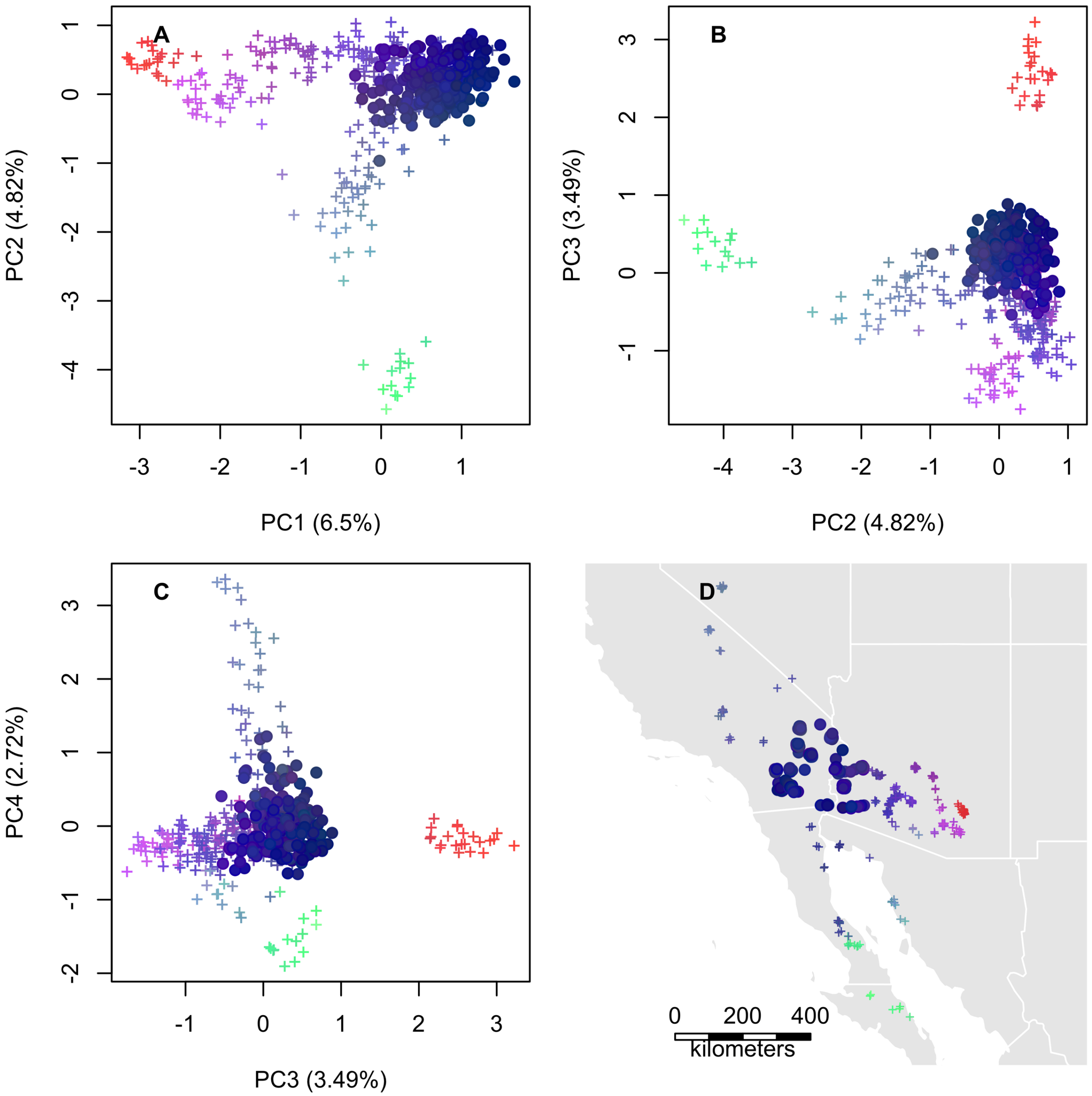


**Figure S1.2.** Principal components analysis of 482 desert clickers based on 1,565 SNPs. **A-C.** Pairwise plots of principal components scores, with points colored by values on PCs 1-3. Individuals selected for tests of genotype-phenotype associations are plotted as circles ($N = 236$). Percentage of variance explained by each PC given in parentheses. **D.** Collection localities for 482 desert clickers (points jittered slightly to reveal multiple samples per site). Plotting colors and symbols as in panels A-C.

**Discriminant analysis of principal components (DAPC)**
We used the find.clusters() function in adegenet to identify the optimal number of genetic clusters. We used all PCs to cluster samples into variable numbers of groups ($K$ = 1-20) using $K$-means clustering, then compared the goodness of fit among results using the Bayesian Information Criterion (BIC). In multiple independent replicates, we found that $K$ = 8 clusters was typically the optimal $K$ (**Fig S1.3**).

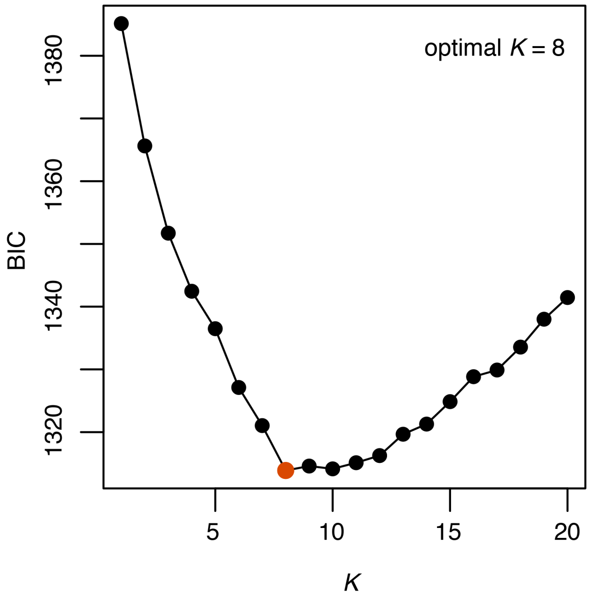


**Figure S1.3.** Comparison of $K$-means clustering results identified $K$ = 8 as the optimal number of genetic clusters.

We then performed discriminant analysis of principal components (DAPC) to probabilistically assign all individuals to one of eight genetic clusters. We used the function optim.a.score() to identify the optimal number of principal components used to construct a discriminant function for cluster assignments. We typically found $N$ = 5-8 PCs as optimal in independent replicates, so we conservatively retained 5 PCs for the final analysis to avoid overfitting the data. Finally, we constructed a discriminant function with dapc() and used all five discriminant axes to assign individuals to clusters.

We visualized the spatial structure of desert clicker populations by mapping the mean assignment probabilities for all sites (**Fig S1.4**). Most sites were dominated by a single genetic cluster, and geographic turnover between clusters was often sharp. However, the boundary between two clusters in western Arizona (orange and red in Fig S1.4) was poorly defined, possibly because isolation by distance was not well captured by a model of discrete clustering. We divided sites into eight groups based upon the majority assignment probability at each site, with one exception. A single population in Baja California was assigned to geographically disjunct populations in the upper Sonoran Desert, so we grouped it with adjacent and genetically similar set of populations in central Baja California.

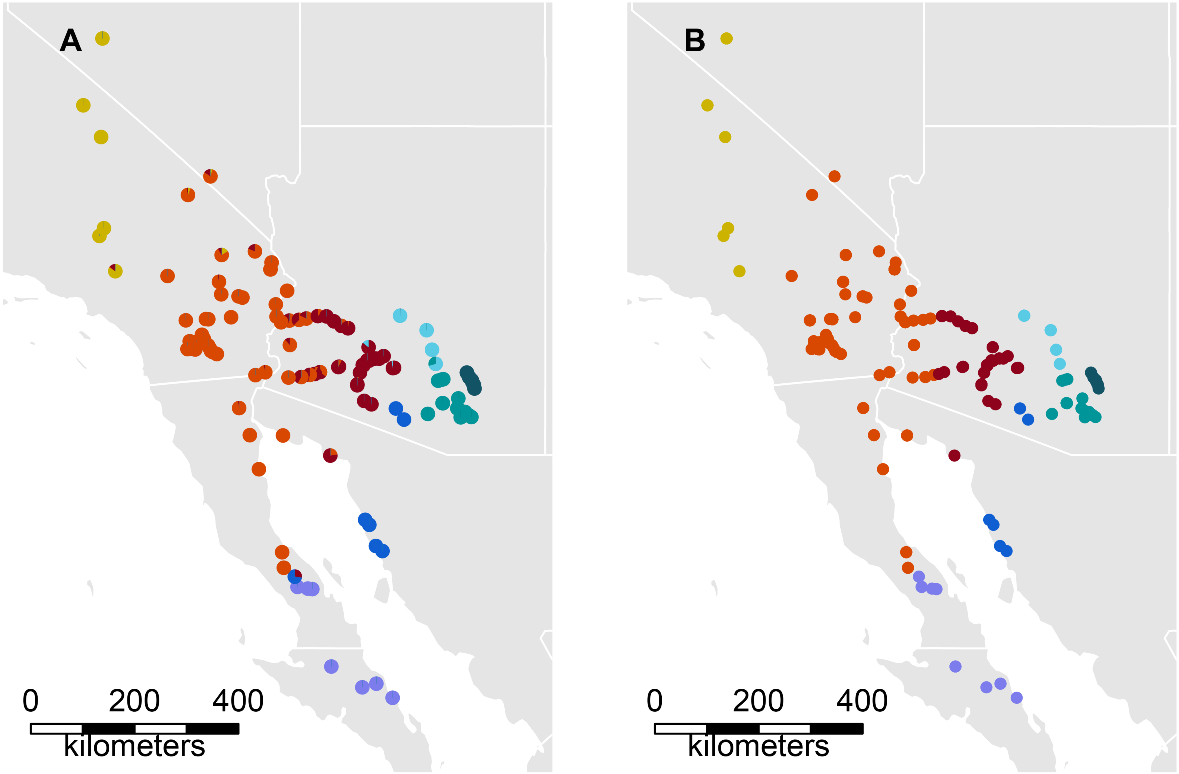


**Figure S1.4. A.** Per-site summary of assignment probabilities from DAPC. **B.** Final population groups based upon DAPC results.

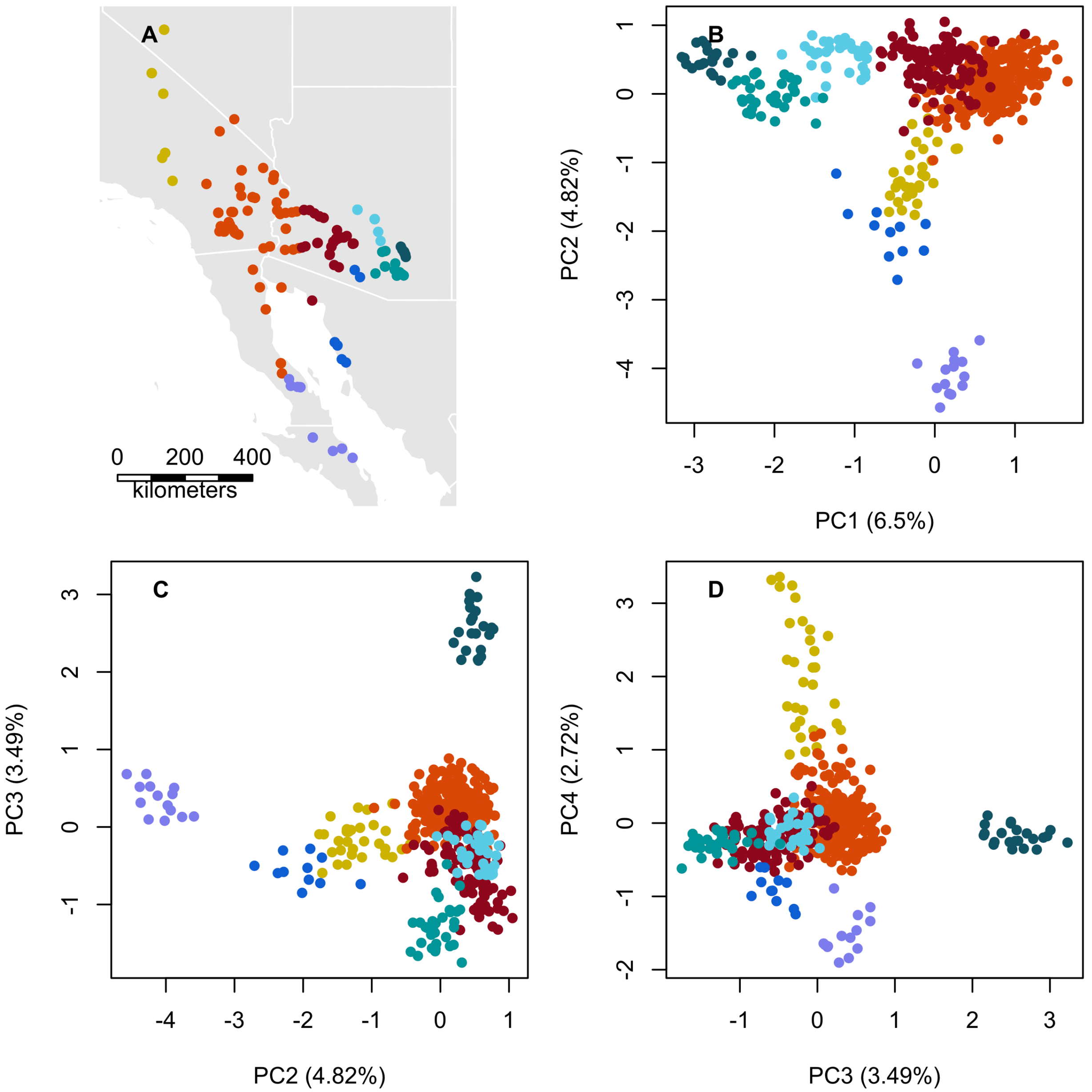


**Figure S1.5.** Summary of genetic variation among desert clicker populations. **A.** Population clusters identified by DAPC (see Fig S1.4 B). **B-D.** Results of principal components analysis of 482 individuals based on 1,581 SNPs.
