## Supplementary material for "Ecological basis and genetic architecture of crypsis polymorphism in the desert clicker grasshopper (*Ligurotettix coquilletti*)": S2. Supplementary Results

### SUPPORTING INFORMATION 2: Supplementary results

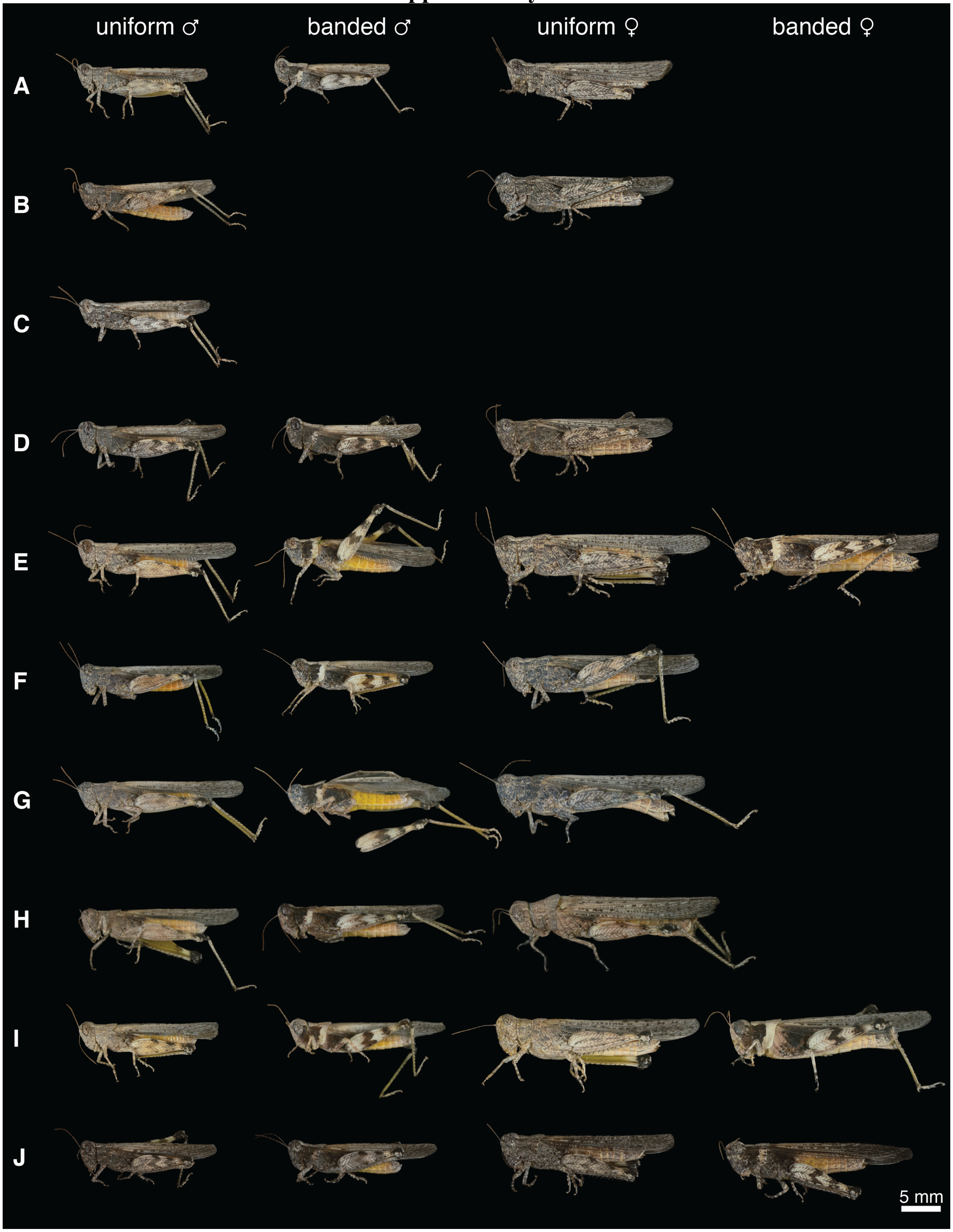


**Figure S2.1.** Typical desert clicker phenotypes at sites A-T. All images are on a common scale and imaged under the same lighting conditions. Note that not all phenotype × sex combinations were observed at every site.

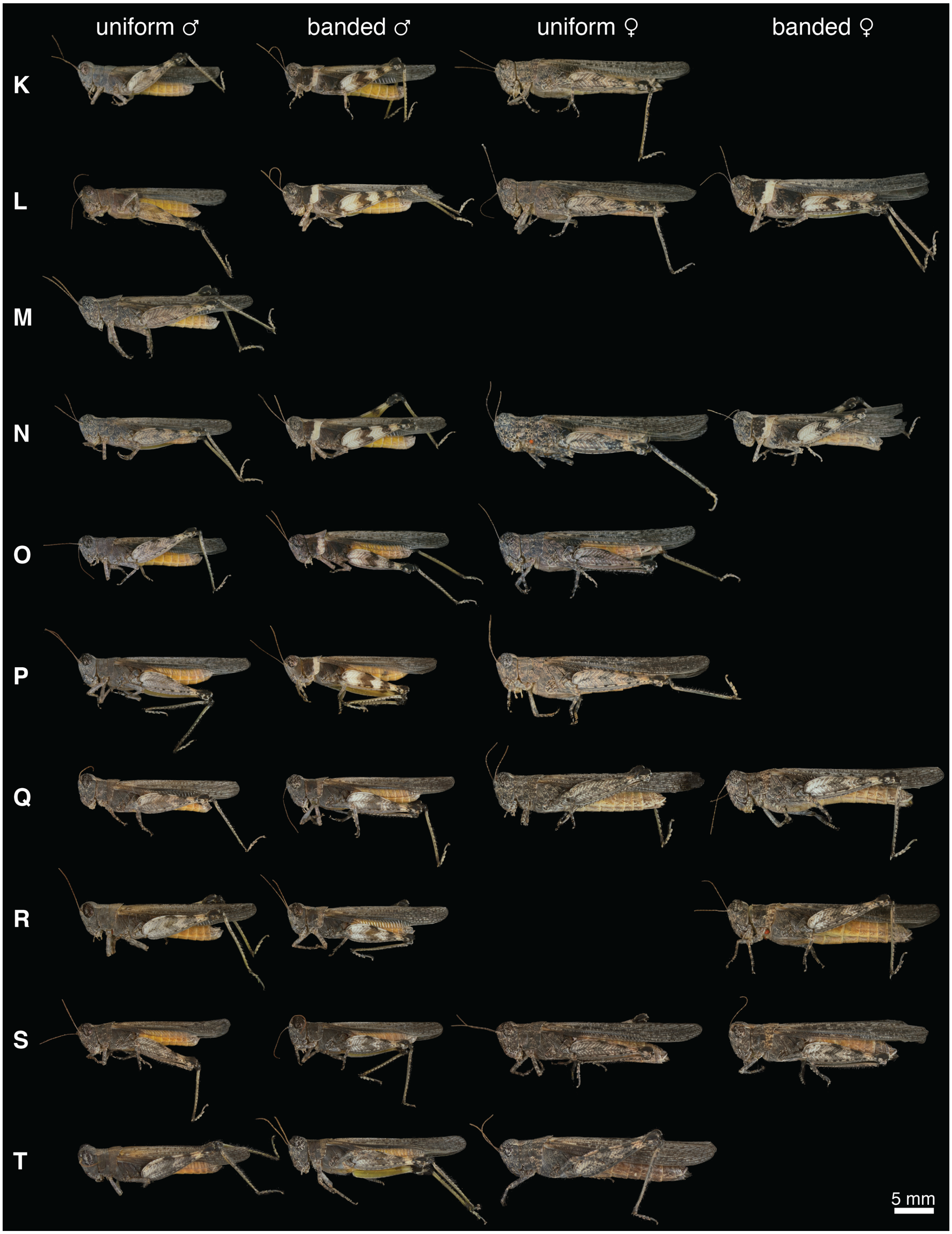

**Figure S2.1** (continued).

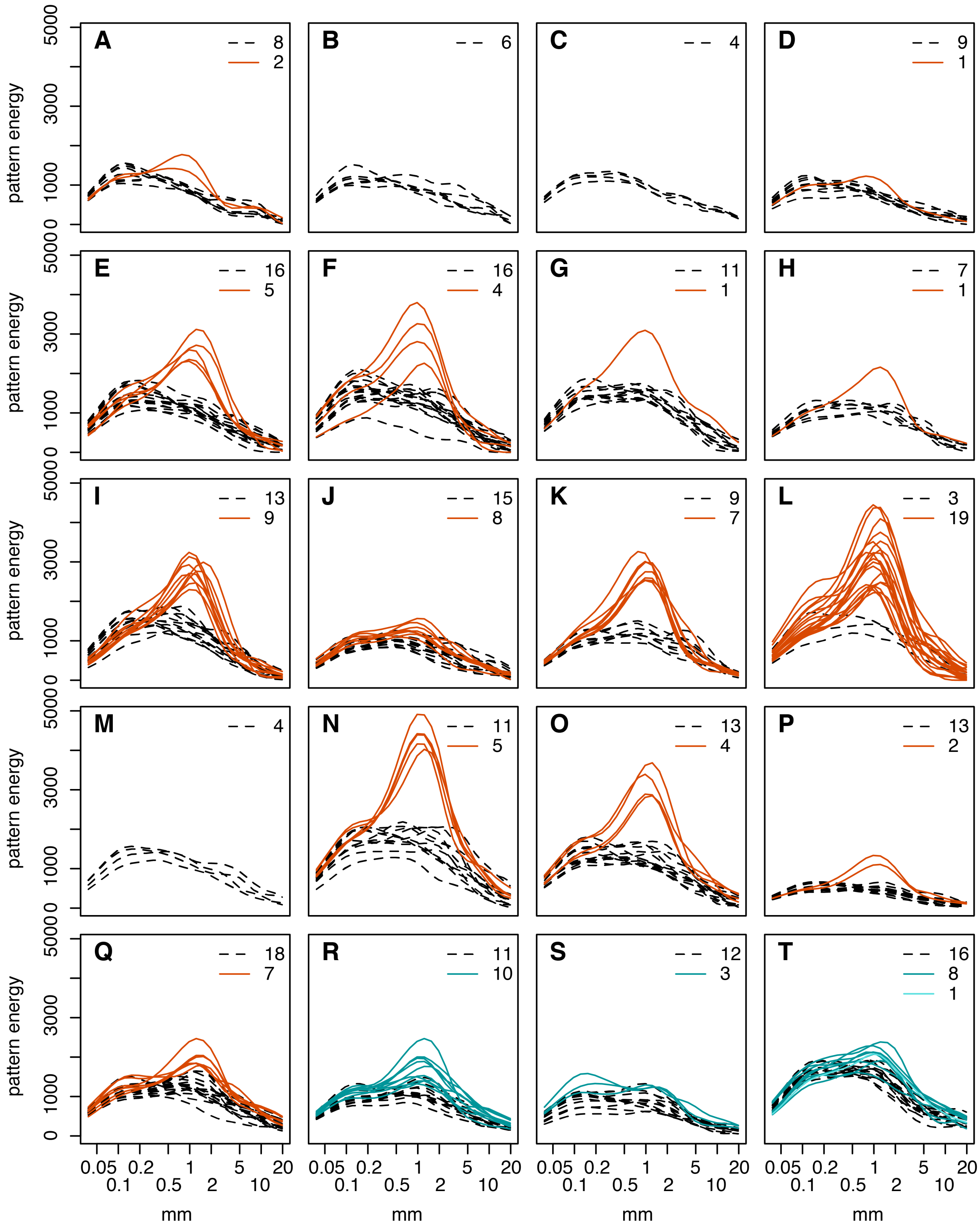


**Figure S2.2.** Comparison of pattern energy spectra between uniform (dashed black) and banded (solid color) grasshoppers at twenty sites (see Fig S1.1 for locations). Numbers of individuals are given in legend. At site T, a single individual of uncertain phenotype is plotted in a lighter color.

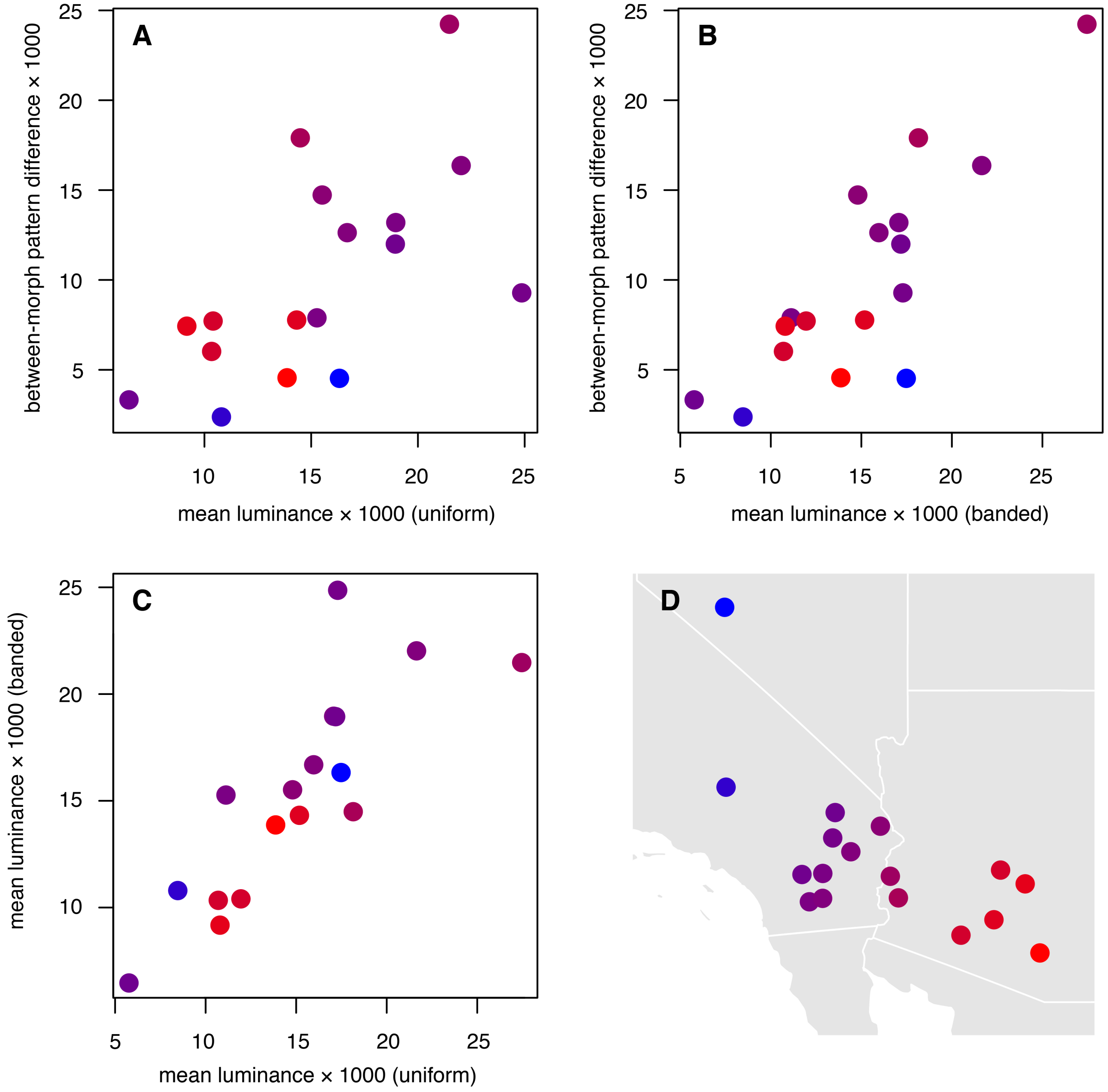


**Figure S2.3.** The difference between the banded and uniform patterns at a given site was predicted by mean luminance of both uniform (**A**) and banded (**B**) grasshoppers. Between-morph pattern difference was calculated as the summed difference between average energy spectra for each morph **C**. Luminance was highly correlated between morphs. **D.** Geographically-based color code used for plotting in panels A-C, highlighting that correlations presented in these panels are not only due to geographic clustering.

**Table S2.1.** Summary of binomial generalized linear model of morph frequencies at 19 sites as a function of substrate patterning (mean PC scores). Significance of model terms was evaluated with a likelihood-ratio $\chi 2$ test. Model $r^2 = 0.60$.

| | ***b*** | **χ2** | **d.f.** | ***P*** |
|---|---|---|---|---|
| substrate PC1 | 0.60 | 29.7 | 1 | **$5.1 \times 10^{-7}$** |
| substrate PC2 | 0.05 | 9.3 | 1 | **$2.3 \times 10^{-3}$** |
| substrate PC1 × substrate PC2 | 0.33 | 7.1 | 1 | **$7.8 \times 10^{-3}$** |

**Table S2.2.** Summary of binomial generalized linear model of morph frequencies at 19 sites as a function of stem patterning (mean PC scores). Significance of model terms was evaluated with a likelihood-ratio $\chi 2$ test. Model $r^2 = 0.02$.

| | ***b*** | **χ2** | **d.f.** | ***P*** |
|---|---|---|---|---|
| stem PC1 | -0.13 | 1.48 | 1 | 0.22 |

**Table S2.3.** Summary of binomial generalized linear model of morph frequencies at 19 sites as a function of substrate and stem patterning (mean PC scores). Significance of model terms was evaluated with a likelihood-ratio $\chi 2$ test. Model $r^2 = 0.63$.

| | ***b*** | **χ2** | **d.f.** | ***P*** |
|---|---|---|---|---|
| substrate PC1 | 0.50 | 21.9 | 1 | **$2.9 \times 10^{-6}$** |
| substrate PC2 | 0.25 | 5.82 | 1 | **0.016** |
| stem PC1 | 0.07 | 0.71 | 1 | 0.40 |
| substrate PC1 × substrate PC2 | 0.22 | 2.27 | 1 | 0.13 |
| substrate PC1 × stem PC1 | -0.11 | 0.51 | 1 | 0.47 |
| substrate PC2 × stem PC1 | -0.34 | 1.72 | 1 | 0.19 |

**Table S2.4.** Summary of binomial generalized linear model of morph frequencies at 16 sites as a function of substrate and stem patterning (mean PC scores). Significance of model terms was evaluated with a likelihood-ratio $\chi 2$ test. Model $r^2 = 0.73$. Model excludes three sites in eastern Arizona where banded and uniform morphs were poorly differentiated. The B haplotype was also found to be recessive at the three excluded sites (see main text for details).

| | ***b*** | **χ2** | **d.f.** | ***P*** |
|---|---|---|---|---|
| substrate PC1 | 0.65 | 24.2 | 1 | **$8.6 \times 10^{-7}$** |
| substrate PC2 | -0.05 | 4.1 | 1 | **0.043** |
| stem PC1 | 0.09 | 0.18 | 1 | 0.67 |
| substrate PC1 × substrate PC2 | 0.43 | 5.56 | 1 | **0.018** |
| substrate PC1 × stem PC1 | -0.14 | 0.64 | 1 | 0.42 |
| substrate PC2 × stem PC1 | -0.04 | 0.02 | 1 | 0.89 |

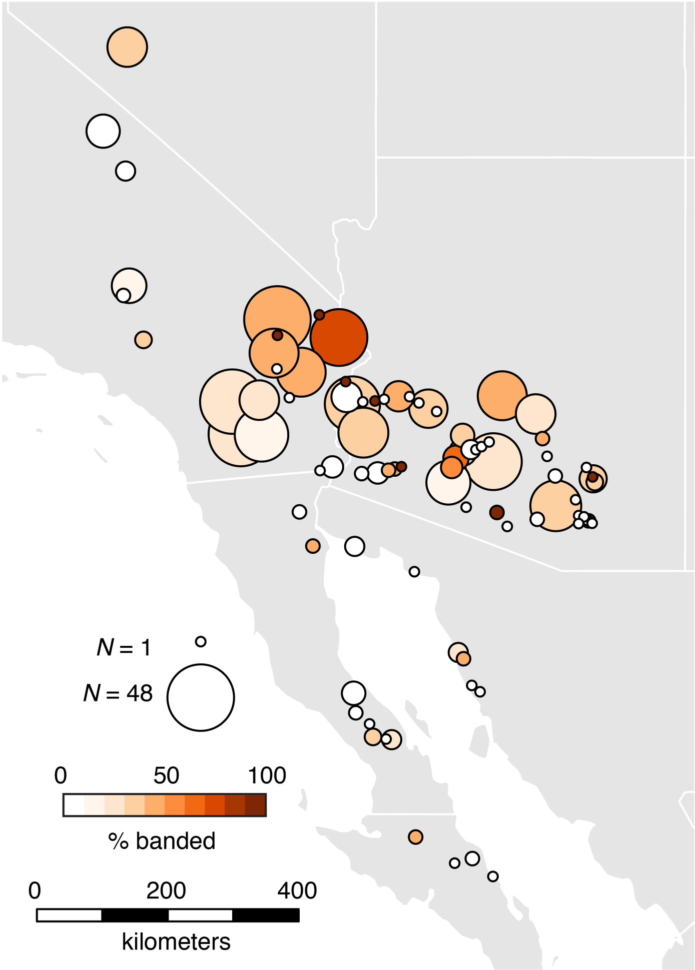


**Figure S2.4.** Local banded morph frequencies summarized from 691 observations across the range of the desert clicker.

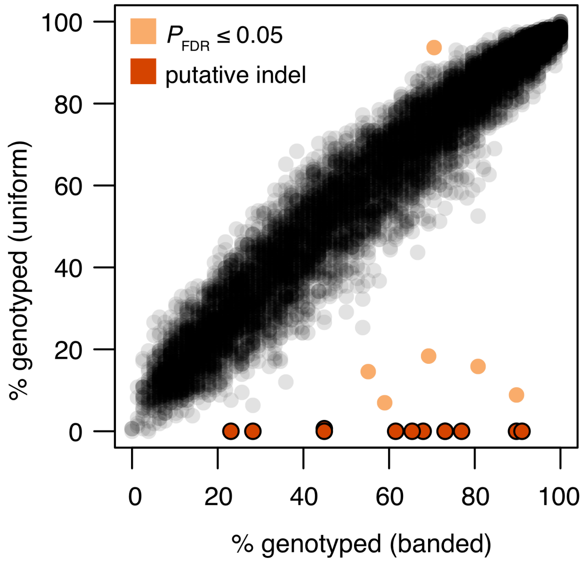


**Figure S2.5.** Comparison of genotyping rates in banded and uniform grasshoppers in western Sonoran populations. Eighteen loci were genotyped at different rates between morphs (Fisher's exact test, $P_{FDR} \leq 0.05$; light orange). Twelve of which were found exclusively in banded morphs (or nearly so), consistent with the presence of a large indel (dark orange).

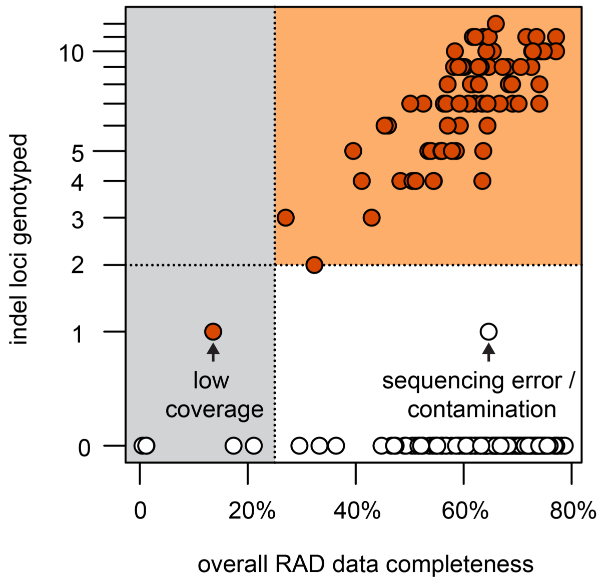


**Figure S2.6.** Rationale for karyotype imputation criteria. Plot shows number of putative indel loci genotyped as a function of overall data completeness for 236 individuals in the western Sonoran Desert (white = uniform, orange = banded). Indel loci were exclusively genotyped in banded individuals (with one exception) and the number of indel loci genotyped was strongly correlated with overall data completeness ($r^2$ = 0.66). Together, these findings indicate that the presence of any indel loci is sufficient to identify an individual carrying a B haplotype. However, we observed one uniform individual genotyped at a single indel locus. Sequencing errors such as index hopping (Illumina 2021) or cross-sample contamination can occasionally cause a genotype to be spuriously associated with the wrong individual. Indeed, we observed a low level of index hopping in our data (e.g., three alleles for a single individual, with one allele being sequenced at much lower depth). A possible example of index hopping is shown in the lower-right. We therefore required an individual to be genotyped ≥ 2 indel loci to impute a B- karyotype. At this threshold, low data completeness can cause some individuals that are likely to carry a B haplotype to be imputed with a UU karyotype (a likely example is shown in lower left). All banded individuals with ≥ 25% data completeness were genotyped at least two indel loci, so we applied this completeness threshold for inclusion in further analyses. Individuals in the gray region were excluded from analysis, individuals in the white region were scored as UU, and individuals in the orange region were scored as B-.

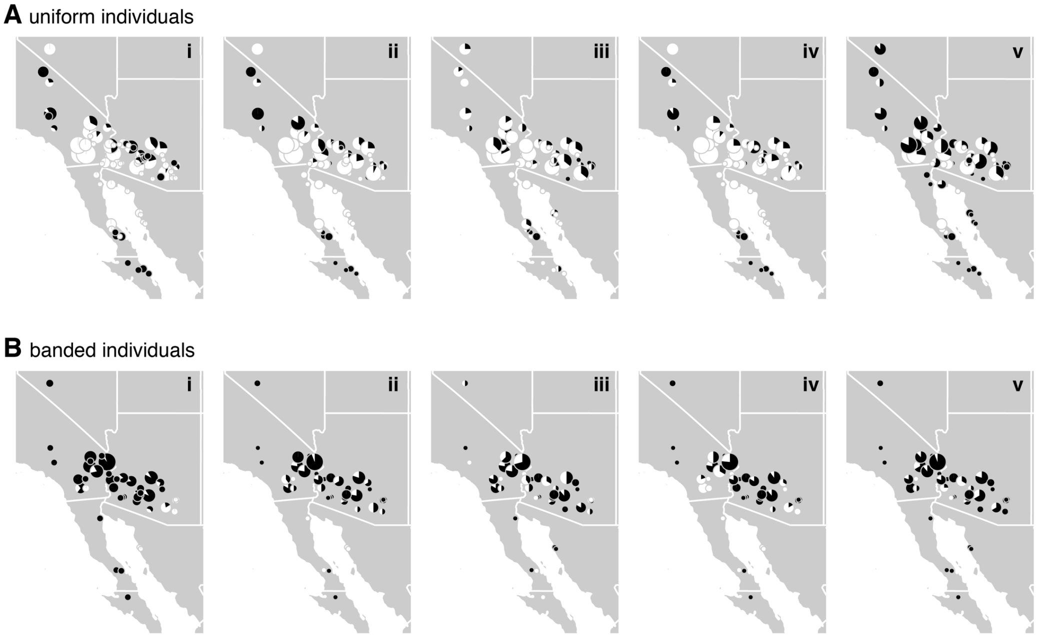


**Figure S2.7.** Geographic patterns of genotyping success for five loci overrepresented in banded morphs (see Fig S2.5). **A.** Uniform individuals. **B.** Banded individuals. In both panels, circle size is proportional to the number of individuals and pies show proportion of individuals that were (black) or were not (white) genotyped at each of five loci (i-v). Genotyping success for four loci (i-iv) show clear geographic signal in uniform individuals, with few individuals from the western Sonoran Desert and northern Peninsular Desert successfully genotyped. This suggests linkage between a cut-site polymorphism and the locus underlying color pattern in those populations.

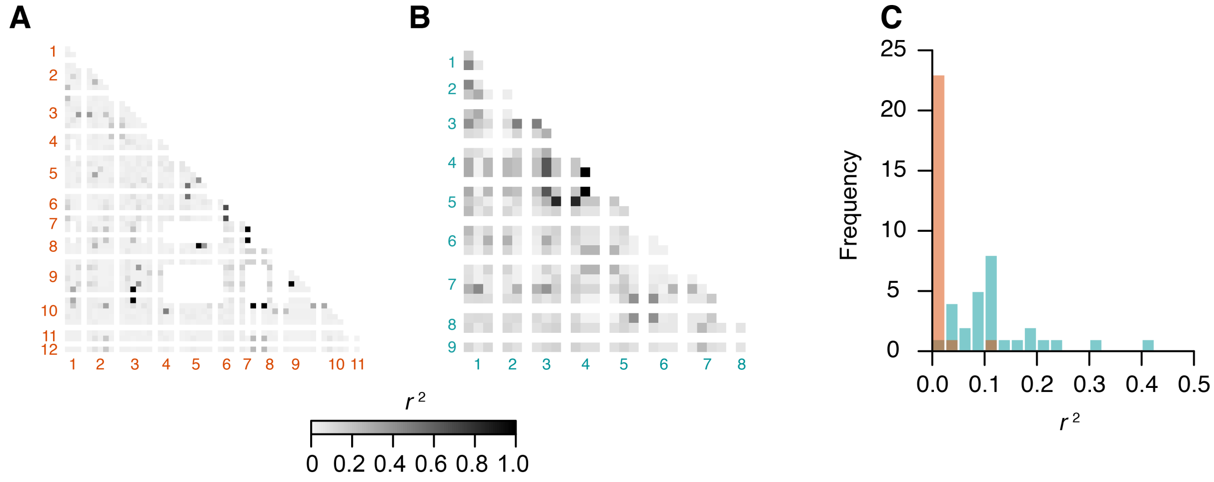


**Figure S2.8.** Patterns of LD between SNPs in 12 indel loci. **A**. Populations in which the B haplotype is dominant (*N* = 109 individuals). Locus numbers indicated at margins of heatmap. Rows / columns = individual SNPs. White = missing data (pairwise genotypes available for <20% of individuals). **B.** Populations in which the B haplotype is recessive in eastern Arizona (*N* = 38 individuals). Due to limited genotyping success, three loci were unavailable for LD calculations (loci 10-12). **C.** Histogram of median between-locus LD (genotypic $r^2$) for both populations. Orange = B dominant, blue-green = B recessive in eastern Arizona. Histogram only includes loci 1-9 to allow direct comparison between populations.

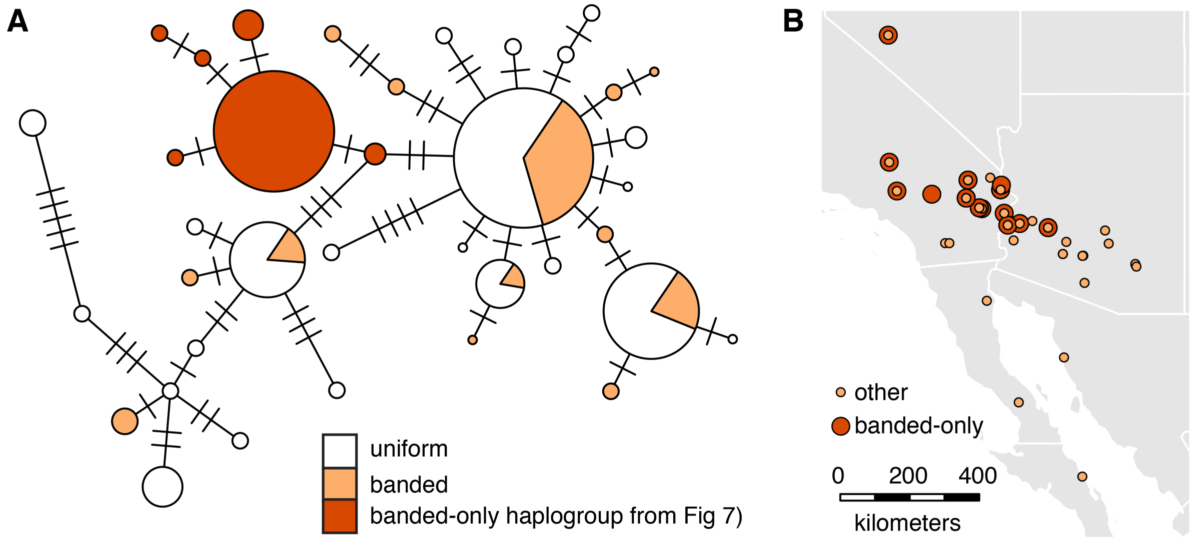


**Figure S2.9. A.** Haplotype network of the morph-associated locus from across the desert clicker's range. Circle size is proportional to number of haplotypes in a haplogroup (max = 75, min = 1). Mutational steps are marked with hashes. The haplogroup found only in banded individuals is shown in dark orange. **B.** Geographic distribution of haplotypes from banded individuals presented in panel A. Haplotypes from the banded-only haplogroup are geographically restricted to the northwestern portion of the desert clicker's range, indicating linkage between the morph-associated locus and locus underlying color morph is incomplete.

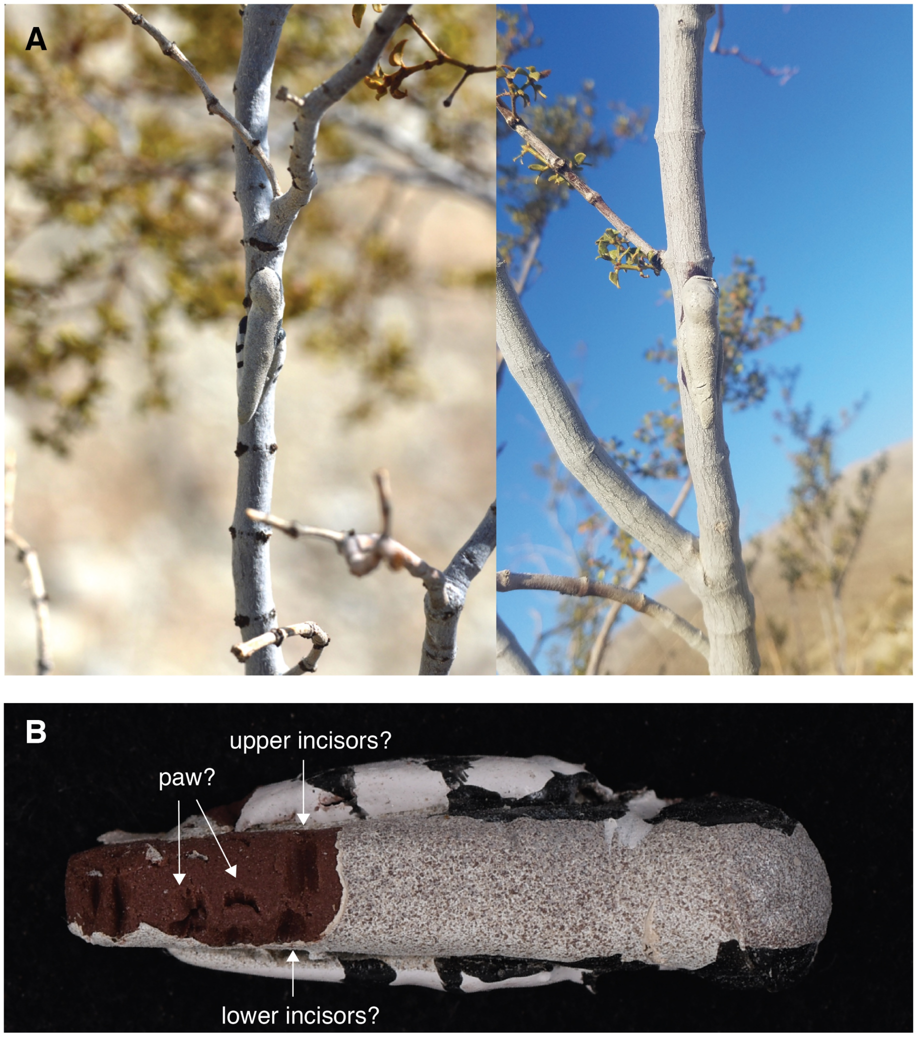


**Figure S2.10. A.** In a small pilot project, we deployed 420 hand-painted plasticine models (210 of each morph) in naturalistic positions on creosote bush (*Larrea tridentata*) at the Boyd Deep Canyon Desert Research Center near Palm Desert, CA. At the end of six days, one model of each morph had inconclusive evidence of predator attacks (markings could not reliably be distinguished from damage due to contact with swaying plant stems). Many confirmed predators of desert clickers (robber flies, spiders, mantids, *Capnobotes* katydids, as well as *Uta* and *Cnemidophorus* lizards) rely upon movement to detect prey and may not have been attracted to models. **B.** Ten models were left on the ground at the base of a host plant for one night. One of these showed clear evidence of predation, including apparent incisor and paw markings consistent with a rodent attack. A grasshopper mouse (*Onychomys*) is a plausible culprit.

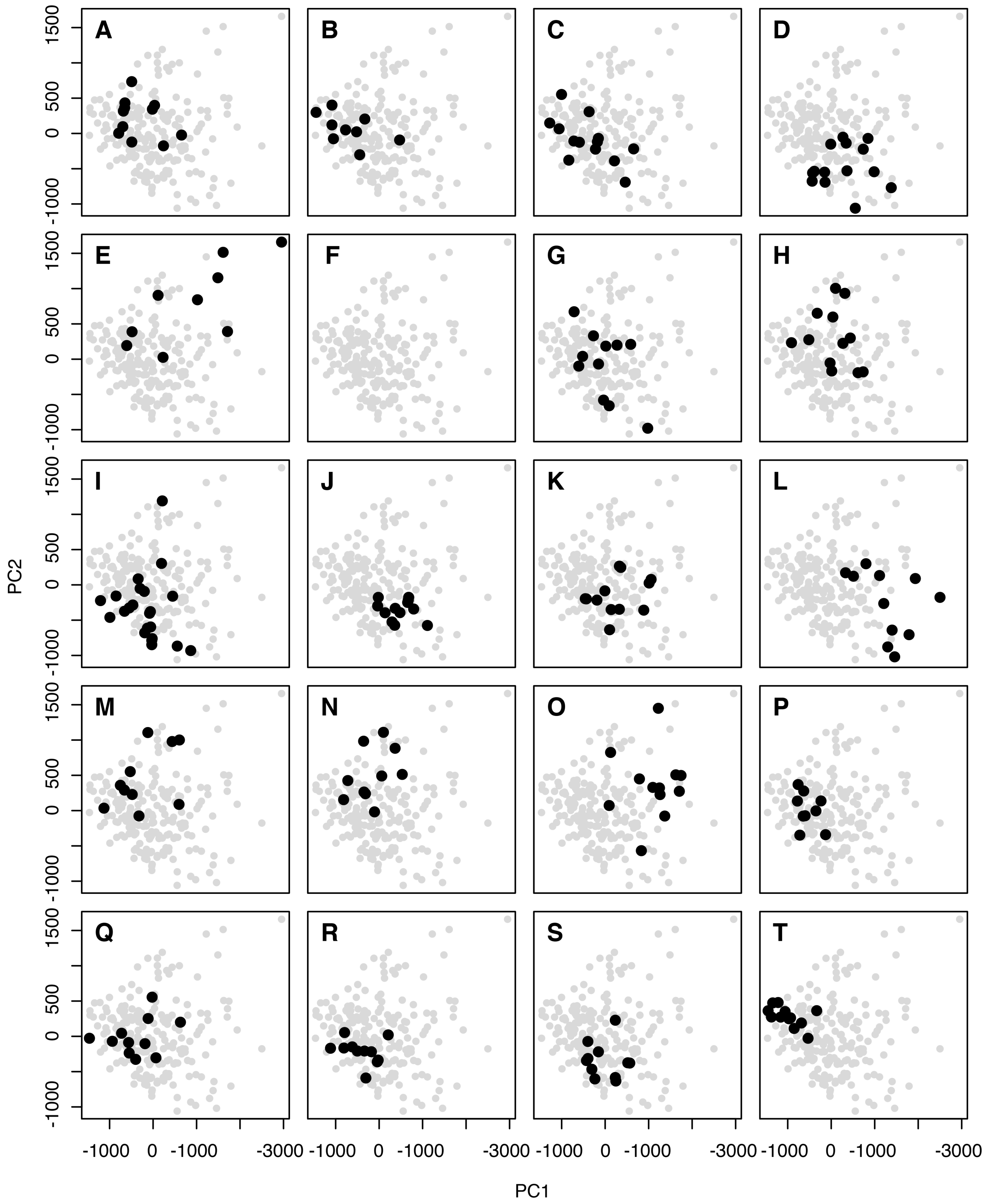


**Figure S2.11.** Principal components analysis of substrate patterning. Panels A-T show scores on PC1 and PC2 for each site in black (see Fig S1.1 for locations) overlaid upon scores for all images in gray. There is no evidence of bimodal substrate patterning within sites that might constitute alternative patches in a Levene model of spatially varying selection. Substrate images were unavailable for site F.

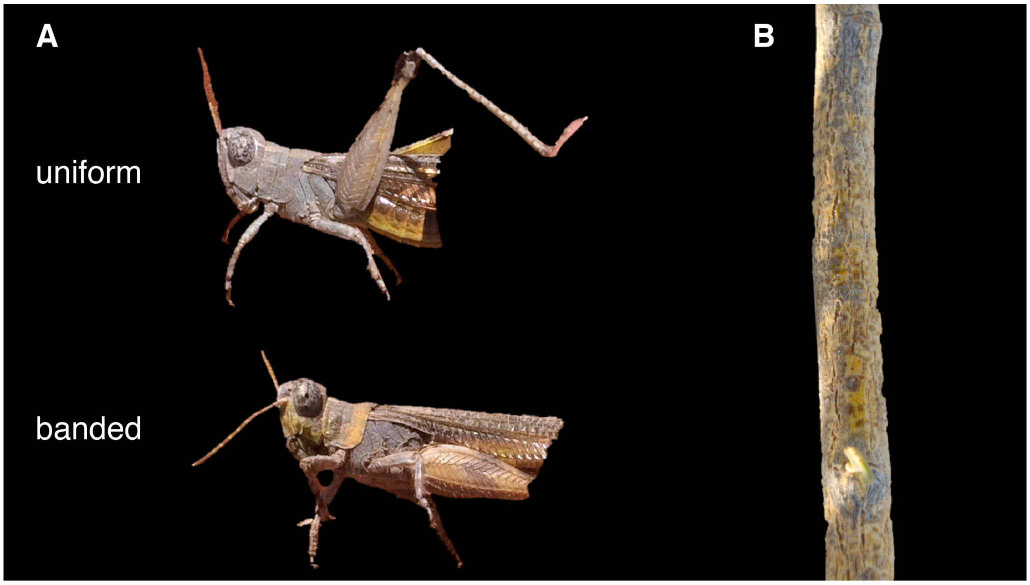


**Figure S2.12. A.** An undescribed species of *Ligurotettix* from Baja California shows a similar color polymorphism as *L. coquilletti*. Top: uniform. Bottom: banded. While coloration differs in banded individuals (gray and yellow instead of black and white), the pattern of different color elements is similar. **B**. Stem of *Viscainoa geniculata*, the primary host plant for the undescribed species.
